# Capacitance measurements for assessing DNA origami nanostructures

**DOI:** 10.1101/2023.03.02.530881

**Authors:** Vismaya Walawalkar, Md. Sakibur Sajal, Yann Gilpin, Marc Dandin, Rebecca E. Taylor

**Author notes:** equal contribution (Vismaya Walawalkar, Md. Sakibur Sajal).

## Abstract

Nanostructures fabricated with DNA are emerging as a practical approach for applications ranging from advanced manufacturing to therapeutics. To support the strides made in improving accessibility and facilitating commercialization of DNA nanostructure applications, we identify the need for a rapid characterization approach that aids nanostructure production. In our work, we introduce a low-fidelity characterization approach that provides an interdependent assessment of DNA origami formation, concentration and morphology using capacitance sensing. Change in charge is one of the transduction methods to determine capacitive loading on a substrate. It is known that cations in the solution stabilize DNA origami nanostructures. So, we hypothesized that the presence of cations and nanostructures in a buffer solution can induce capacitance change that is distinctive of the nanostructure present. In this study we were able to detect a change in the capacitance when the nanostructure solution was deposited on our capacitance sensor, and we could distinguish between pre-annealed and annealed structures at concentrations less than 15 nM. The capacitance measurements were affected by the concentration of Mg^2+^ ions in the solution, the staple-to-scaffold stoichiometric ratio of the nanostructure and the nanostructure morphology. Maintaining a 12.5 mM Mg^2+^ concentration in the nanostructure buffer, we discover a linear relationship between the relative capacitance change and the nanostructure concentration from 5 nM to 20 nM, which we call the characteristic curve. We find distinct characteristic curves for our three nanostructures with distinct morphologies but similar molecular weight - a rectangular plate, a sphere and a rod. Given that we can distinguish nanostructure formation, concentration and morphology, we expect that capacitance measurement will emerge as an affordable and rapid approach for quality control for nanostructure production.

## Introduction

Building nanostructures^1–3^ from nucleic acids enables addressable placement of moieties^4–6^ and dynamic responsive behavior. ^7–9^ Given these attributes, DNA nanostructures can be excellent candidates for applications such as molecular sensing and actuation, ^10–14^ creating organic-inorganic hybrid materials^15,16^ as well as in physiological studies^9,17–19^ and therapeutics.^20,21^ A leading approach in structural DNA nanotechnology is DNA origami, in which nanostructures can be self-assembled at high yield into various architectures including honey-comb lattice, square lattice or wireframe. ^2,22,23^ The common scheme for DNA origami self-assembly uses a long circular single strand DNA called ‘scaffold’ and a 5-fold or 10-fold excess of unique short single-stranded DNA (ssDNA) strands called ‘staples’.^22,24^ Self-assembly yields for these nanostructures can be as high as 70 - 90%. The result of nanostructure synthesis includes well-formed DNA origami and excess staples, unbound scaffold, partially-folded structures or aggregated structures.

To verify that a large portion of well-formed product is present, various methods from analytical chemistry and microscopy are typically utilized to assess formation and morphology, purify samples and determine yield. ^24,25^ Researchers typically utilize multiple complementary methods to evaluate their nanostructures.^25^ These techniques tend to fall into two categories: (1) high fidelity characterization approaches are essential for quantitative validation of nanostructure dimensions and morphologies and (2) low fidelity approaches that are essential for qualitative or semi-quantitative assessment of formation at the bulk solution level.

Various characterization techniques require different levels of complexity in terms of infrastructure and accessibility and provide a range of information in terms of quantification (Figure 1). High fidelity approaches tend to be highly complex and provide quantitative measurements. Users typically select one or more high fidelity approaches based on their strengths and weaknesses. For example, standard atomic force microscopy (AFM) provides a high resolution visualization of the morphology and mechanics of nanostructures,^26–28^ but deposition on mica can lead to undesirable deformation and loss of structural integrity.^29^ Transmission electron microscopy (TEM) provides a high resolution projected view of stained or unstained nanostructures,^30–32^ which has been critical for visualizing delicate and curved nanostructures, with the potential for high throughput imaging to generate 2D class average images.^32^ Cryo-electron microscopy (cryo-EM) enables structural determination with °Angstrom-scale resolution^33^ and even construction of pseudoatomic models.^34^ Emerging high fidelity approaches like DNA points accumulation for imaging in nanoscale topography (DNA-PAINT) enable more traditional fluorescence microscopy approaches to achieve super resolution with steep reductions in equipment costs.^35^ Approaches such as small-angle x-ray scattering (SAXS) can enable nanostructure distance measurements with resolution as low as 1.2 nm.^36^ Although the complexity of SAXS experiments is high, SAXS generates a unique signature for a each morphology ^37^ and can also be performed dynamically during a dilution.^38^

**Figure 1:**
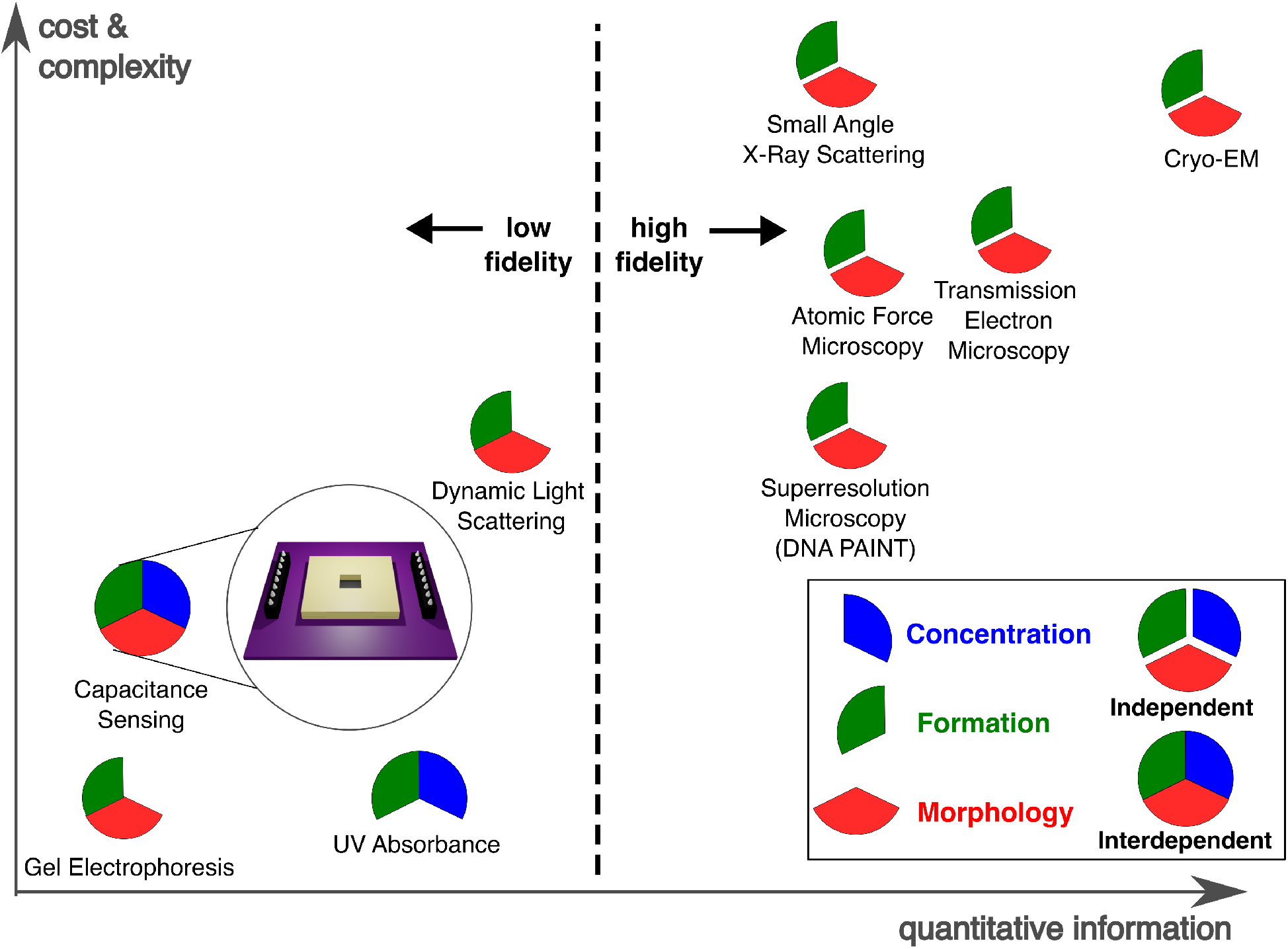
Formation, concentration and morphology form the basis of nanostructure assessment. Although high fidelity assessments deliver independent and quantitative information of the nanostructures, they are generally higher in cost and complexity. Low fidelity nanostructure characterization approaches deliver interdependent assessment of the nanostructures, at low cost and complexity, and are more suitable for routine nanostructure assessments.

While emerging innovations and automation are bringing down the cost and complexity of high fidelity measurements, the field needs fast and low-cost characterization methods for daily use in the lab or for production. Low fidelity measurements like gel electrophoresis, dynamic light scattering (DLS) and UV absorbance provide a more practical assessment of nanostructure formation, concentration and morphology. Unlike high-fidelity approaches that allow independent assessment of these criteria, low-fidelity measurement provides an interdependent signal that describes the interaction between multiple variables. For example, UV absorbance rapidly provides a measure of concentration under the assumption of a well-formed structure.^39,40^ Gel electrophoresis is a low cost but slow assay that can distinguish between different morphologies of nanostructures with the same molecular weight. Gel band intensity can also be related to yield or concentration, while the spread or smearing of a band can speak with formation and degradation.^1^ Similarly, recent work on DLS has shown that in addition to providing particle size distributions, DLS can be used to monitor and characterize changes in morphology due to melting and formation as well as degradation events.^41^ Importantly, while DLS has emerging capabilities and is amenable to high-throughput measurement, the high instrument cost may limit its accessibility.

Consequently, we recognize the need for low fidelity approaches that are both rapid and low cost. Such an approach would not only improve the accessibility of DNA nanotechnology, but it would also facilitate commercialization efforts by enabling enhanced quality control of nanostructure production. Here we present an affordable and rapid approach that uses CMOS-based capacitance sensing for DNA origami assessment. We anticipate that our approach will provide a potential high-throughput, and low-effort alternative to gel electrophoresis for assessment of validated nanostructures.

We show that the presence of DNA origami nanostructures in a solution elicits interdependent changes in capacitance that correlate with the formation, concentration and morphology of DNA origami nanostructures. To establish the feasibility of such detection, we investigate the impact of staple stoichiometry, Mg^2+^ concentration, and origami concentration on capacitance change. The capacitance changes are read via an array of paired electrodes within a lab-on-CMOS system. The system transduces the presence of charged particles, like DNA origami, as a change in capacitance at each electrode pair. ^42^ The negative backbone of the DNA origami nanostructures are often stabilized by adding salts of Mg^2+^ or Na^+^ to a buffer with a low pH.^43^ So, the capacitance of the nanostructure in solution can be affected by both the nanostructure and the solution constituents. Therefore, we introduce a relative capacitance metric that isolates the effect of the nanostructure on the capacitive signal. In addition, since our capacitance sensor is primarily designed to detect cell viability, ^44^ our relative capacitance metric adapts the system for assessment of DNA nanostructures.

After demonstrating the dependence of the relative capacitance on Mg^2+^ concentration, we investigate the utility of this low-fidelity approach for assessing formation, concentration and morphology of DNA nanostructures at fixed concentrations of Mg^2+^. We find that relative capacitance is approximately linear with nanostructure concentration, and that interdependent capacitive measurements across a range of nanostructures can be used to successfully distinguish pre-anneal versus post-anneal nanostructures as well as nanostructure morphologies.

## Results and discussion

We study capacitive response to the formation, concentration and morphology of three DNA origami nanostructures. Each nanostructure has similar molecular weight, being assembled with M13mp18 scaffold and about 200 unique staples. The nanostructure sequences are listed in Supplementary Information. We regard nanostructure morphology to be a combination of the architecture, dimensions and geometry. Each nanostructure has a distinct morphology: a planar square-lattice nanotile (74 nm × 96 nm), a wireframe sphere (diameter = 51 nm) and a honeycomb-lattice rod (425 nm × 7 nm). We confirm the formation of the nanostructures and obtain the dimensions using atomic force microscopy (AFM) (Figure 2). We deposit each of these nanostructures on our lab-on-chip capacitive sensing platform to obtain their distinctive capacitance signatures.

**Figure 2:**
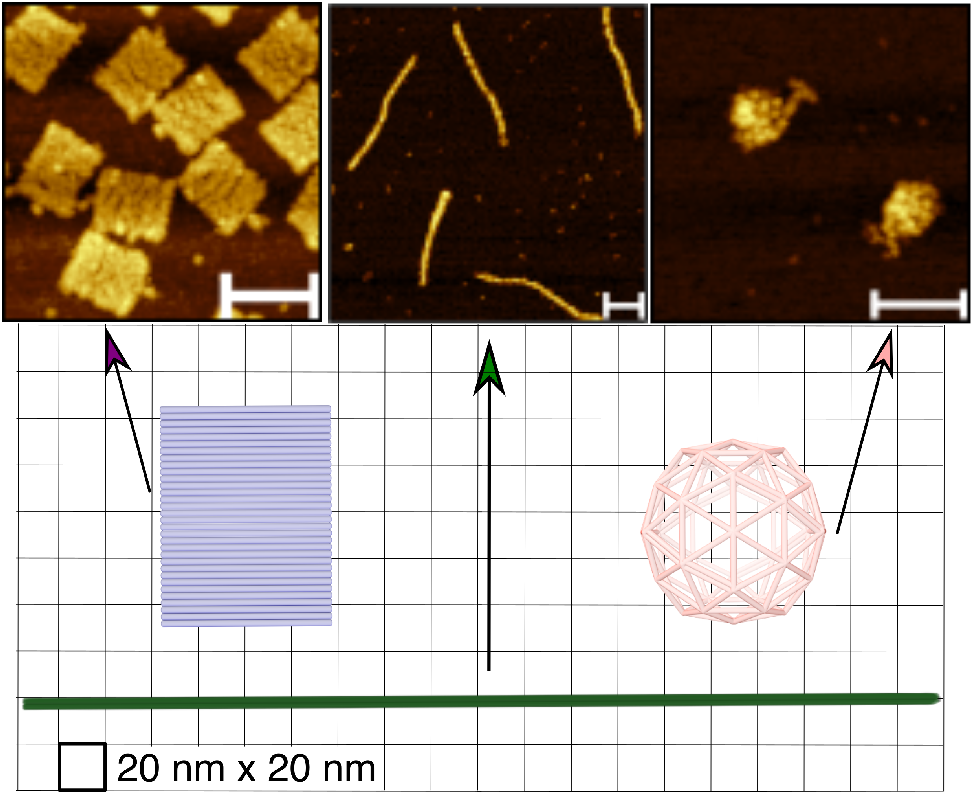
Atomic force microscopy characterization and scaled schematics for three distinct DNA origami nanostructures: a nanotile, a nanorod, and a nanosphere. Scale bars: 100 nm.

### Relative capacitance change (r∆C) varies with formation, concentration, and morphology of DNA nanostructures

The distribution of the ions in the solution containing the DNA origami nanostructures relative to the nanostructures’ negatively charged backbones likely gives rise to the capacitive effects measured by our platform’s sensing electrodes. Specifically, because nanostructures have negatively charged backbones and are often stabilized with ionic salts such as MgCl_2_, and in buffers such as 1xTAE (40 nM Tris base, 20 mM acetic acid, 1 mM EDTA),^45^ the application of an electric field by a set of electrodes to this system is expected to induce dipoles at the electrodes’ interface, which changes the electrodes’ capacitance (Figure 3).

**Figure 3:**
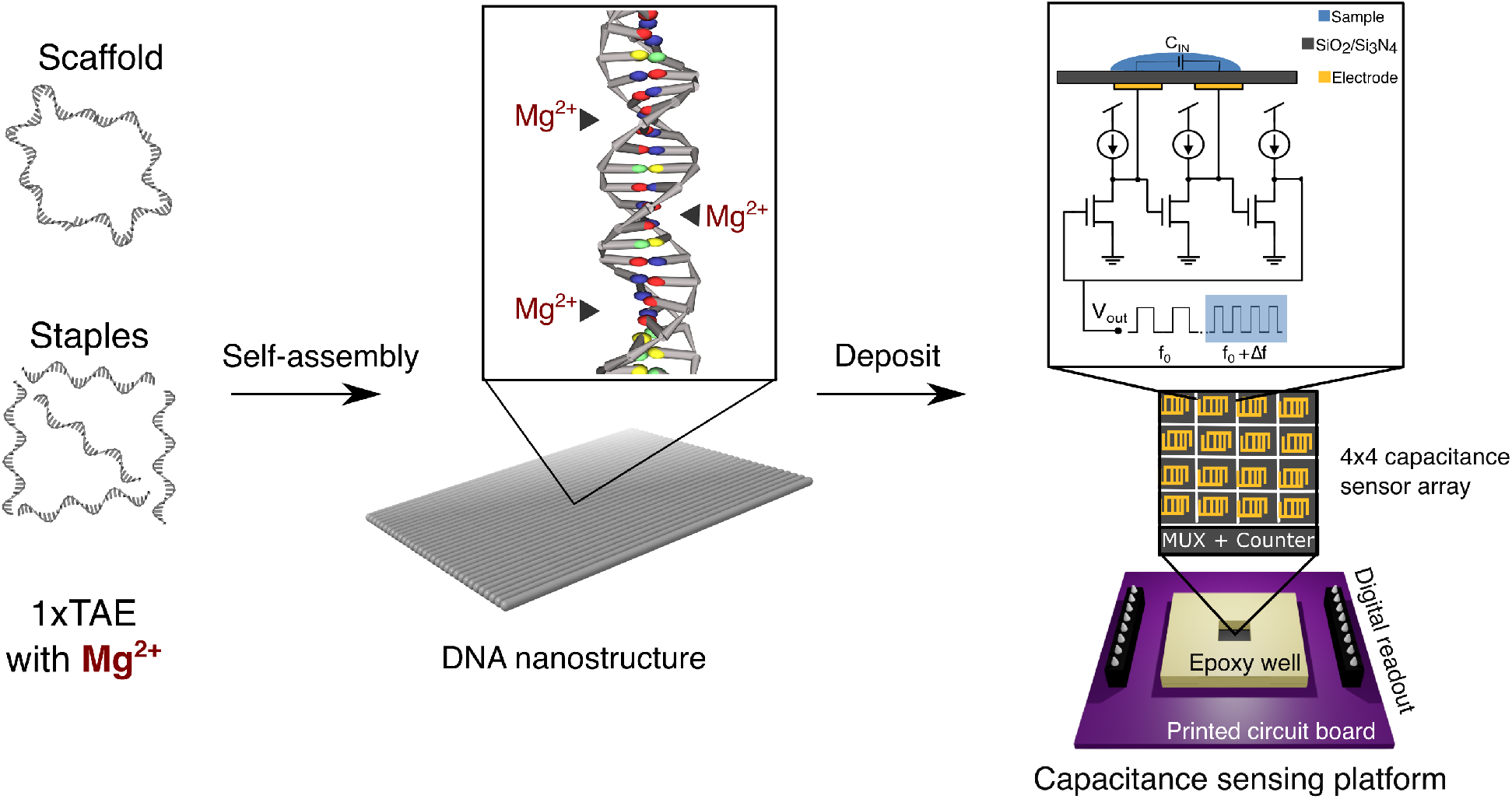
Assessing formation, concentration, and morphology of DNA nanostructures is possible by measuring relative capacitance noted in Eq. (3) based on the concentration of the DNA nanostructures and Mg^2+^ ions in a solution. (left) Self-assembly of DNA scaffold and staples in a 1xTAE buffer with Mg^2+^ (middle) forms DNA nanostructures, (right) which effect a change in the output voltage frequency. Relative capacitance (r∆C) is calculated from the number of oscillations in a given integration time.

### Capacitance sensing platform

Our capacitance sensing platform is an integrated circuit chip fabricated in a 0.35 *µ*m, 4-metal, 2-poly complementary metal-oxide semiconductor (CMOS) process. The chip measures 3 mm × 3 mm, and it integrates 16 capacitance sensors arranged in a 4 × 4 array. Each sensor has a sensing area of 30 *µ*m × 30 *µ*m, and the vertical and horizontal pitches between the sensors are 196 *µ*m and 186 *µ*m, respectively. The chip also includes an I^2^C core for serial communication.

A sensor consists of a pair of interdigitated electrodes disposed under the chip’s top oxynitride passivation layer. The electrode pair is coupled to a dedicated sensing circuit. The presence of DNA origami at the electrodes’ interface is transduced to an electrical signal via a capacitance-to-frequency function implemented by the sensing circuit.

In each sensing circuit, the electrode pair is connected to the intermediate stage of a three-stage NMOS ring oscillator. The oscillator’s base frequency is established by the electrode pair’s intrinsic capacitance. Further, the fringing electric field of this capacitive system extends in the positive z-direction, beyond the passivation layer. Therefore, when the DNA origami solution is applied to the surface of the chip, the fringing field perturbs the charge distribution at the surface, separating positive counterions in the medium and the negatively-charged DNA origami backbones, thereby inducing a plurality of dipoles. These induced dipoles cause a change in the capacitance of the electrode pair, ^42^ which in turn changes the ring oscillator’s frequency. The change in frequency in the test signal, *i*.*e*., the relative change in frequency of the oscillator’s output signal, is estimated by counting the signal’s edges using a digital counter for a fixed integration period. The measured count is routed off-chip via the I^2^C core to a control program that converts it to a relative capacitance measure.^44^

In our experiments, the deposition of nanostructures on the capacitive sensor elicit a capacitive load (C_*IN*_) that maps to the oscillator frequency as *f* (*C*_*IN*_) = *f*_0_ + ∆*f*, where *f*_0_ is the oscillator baseline frequency without the origami and ∆*f* is the change in frequency caused by the presence of the origami over the sensor.

Further, the capacitance *C*_*IN*_ is in parallel with a deterministic stray capacitance *C*_0_ formed by the electrode’s baseline capacitance and the parasitic capacitances of the CMOS stack. Thus, for small changes in capacitance, the capacitance sensor frequency response can be modeled linearly with the estimated capacitance change as shown in Eq. (1),^44^ where *α* is a sensor gain factor (in *Hz/F*) obtained via calibration measurements.

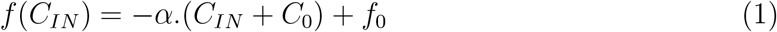

### Relative Capacitance

As illustrated in Supplementary Information (Figure S7), preliminary studies have shown that DNA nanostructures elicit capacitance changes less than 2 fF, and so the sensor output frequency response, f(C_*IN*_), can be modeled linearly as shown in Eq. (1). In Eq. (1), the *α* is the pixel sensitivity that is globally affected by ambient temperature and locally affected the sensor fabrication process. Accordingly, *α* may vary between distinct pixels during a particular trial and also *α* may vary for the same pixel for distinct trials. So, we seek a metric of interest that is independent of *α*. Since *α* responds to ambient conditions, when no sample is deposited, we consider a *pseudo-*load, C_*idle*_, is registered by the sensor. On deposition of a sample solution, we correct for the *pseudo-*load as shown in Eq. (2)

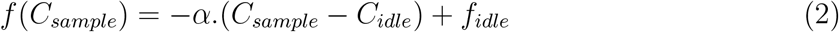

Even so, a sample of DNA origami nanostructures induces a capacitance change as a consequence of the interaction of the negatively charged DNA backbone, the Mg^2+^ and the 1xTAE buffer with pH between 7.5 and 8.2. To isolate the capacitive effect of the sample of interest, we introduce the relative capacitance metric (relative C) as shown in Eq. (3), that corrects and normalizes the capacitive response to deposited sample solution with the capacitive response to corresponding solution without the sample of interest. The derivation for the relative capacitance is detailed in Supplementary Information.

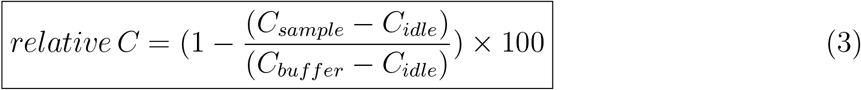

The solution without the sample of interest is the negative control group when studying a given sample. The relative capacitance embeds the negative control measurement to eliminate the need to calibrate our sensor to every DNA nanostructure conditions.

Nevertheless, as the relative capacitance normalizes the data against the solution buffer, we characterize effects of the solution constituents on the capacitance. We expect that the change in Mg^2+^ concentration in the 1xTAE buffer drives the change in relative capacitance.

### Relative capacitance change is influenced by the Mg^2+^concentration

According to Eq. (3), to discern the effect of Mg^2+^ on the capacitance change, we use 1xTAE with no MgCl_2_ as the negative control. We vary Mg^2+^ concentration in 1xTAE from 8 mM to 30 mM, and note the relative capacitance (Figure 4(a)). We use one-way ANOVA test to see the difference between the relative capacitance obtained from different concentrations (*χ*) of Mg^2+^ in 1xTAE. The 95% confidence intervals of the difference between means for 1xTAE with Mg^2+^ concentrations between 8mM Mg^2+^ to 20mM Mg^2+^ are distinct and show no overlap as shown in Table S1 in the Supplementary Information. The 95% confidence intervals of the difference between means, shows that the range of possible relative capacitance for 1xTAE with 20 mM overlaps with that for 30mM Mg^2+^, while the range of possible relative capacitance for 16 mM is overlaps with that for 24mM Mg^2+^ (Figure S8). With these overlaps, we speculate that our capacitance sensor output may begin to saturate near 20 mM of Mg^2+^. Nonetheless, a linear regression fit (0.0714 + 0.504*χ*, R^2^ = 0.889, p-value=0.005) on the relative capacitance for Mg^2+^ concentrations (*χ*) from 8 mM to 30 mM shows that the relative capacitance increases by approximately 0.5 for every 1 mM increase in [Mg^2+^] in 1xTAE (Figure 4(a)).

**Figure 4:**
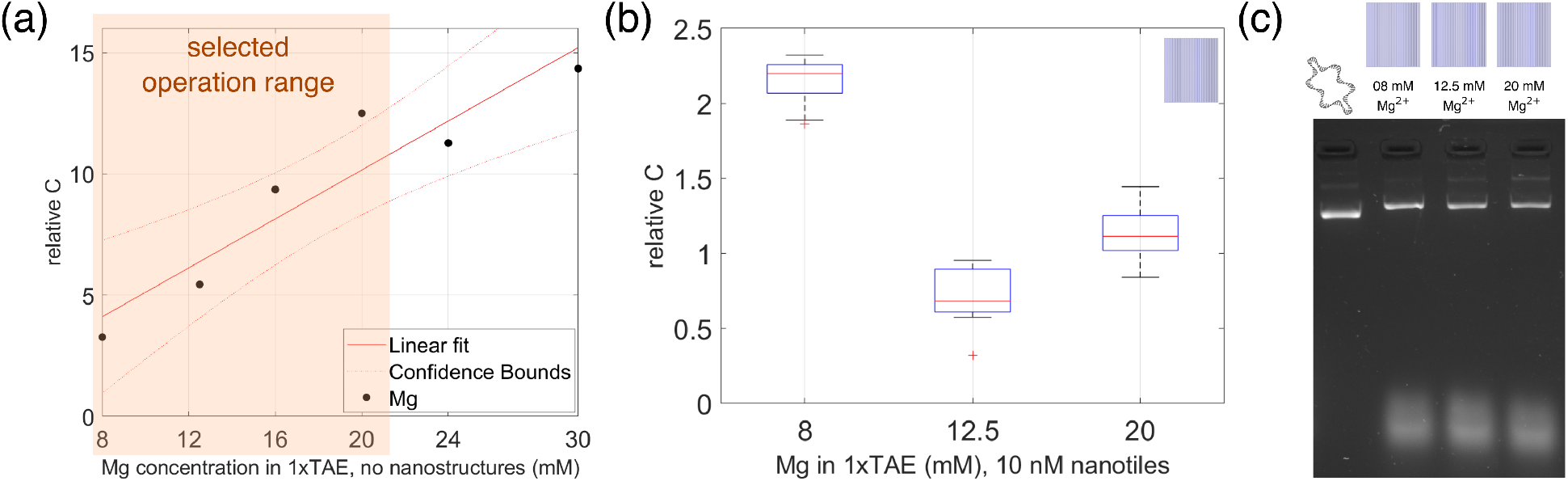
The relative capacitance metric is influenced by the concentration of MgCl_2_ in the DNA origami buffer. (a) A linear approximation (R^2^=0.889 and p-value=0.005) can be fit between the relative capacitance and 8 mM to 30 mM of Mg^2+^ in 1xTAE (N=3, one-way ANOVA p=4.088e-69) containing no nanostructures. The dotted lines show the confidence interval of the linear fit. (b) A nonlinear relationship between the relative capacitance and 10 nM of nanotiles that are annealed and diluted in 1xTAE with 8 mM, 12.5 mM and 20 mM of Mg^2+^ (N=3, one-way ANOVA p=1.114e-26) is illustrated by the box-plot. The red lines inside the box denote the median relative capacitance. (c) Gel electrophoresis image of unpurified nanotiles in 1xTAE with 8 mM, 12.5 mM and 20 mM of Mg^2+^, shows that gel bands for each nanotile variation migrates similarly.

Our linear approximation of the sensor response allows application of Eq. (1). From 8 mM to 20 mM Mg^2+^, we observe a monotonic increase in the relative capacitance, and choose that as our operating range. Our chosen sensor operating range for Mg^2+^ from 8 mM to 20 mM corresponds to the concentration range in literature for successful folding of nanostructures.^25,45,46^ To investigate capacitive signatures of our rectangular planar nanostructures, the nanotiles, we select three Mg^2+^ concentrations in the operating range - 8 mM, 12.5 mM, which represents standard Mg^2+^ concentration for high yield,^46,47^ and 20 mM, which represents the probable saturation band. We speculate that increase in Mg^2+^ concentration in the buffer can desensitize the sensor by masking the small changes in the capacitive measurements that may occur on the sensor surface. We expect that higher Mg^2+^ concentration will note lesser relative capacitance for DNA nanostructures. The nanotiles are annealed separately in 1xTAE buffer with 8 mM, 12.5 mM and 20 mM MgCl_2_ and the negative control is the corresponding annealing buffer and the relative capacitance for 10 nM of unpurified nanotiles is noted for different annealing buffers (Figure 4(b)). The relative capacitance shows a non-linear dependence on Mg^2+^ concentration in the buffer. The primary gel bands for agarose gel electrophoresis of the nanotiles in varying Mg^2+^ concentrations (Figure 4(c)) of 8 mM, 12.5 mM and 20 mM Mg^2+^ are identified by their proximity to the gel band for the M13 scaffold. The primary bands migrate comparable lengths due to similar molecular weights and aspect ratios of all three nanotile conditions, which demonstrates that gel electrophoresis does not note the buffer conditions of the nanostructures, whereas relative capacitance provides distinct readings for each buffer condition. The wider gel bands that travel the furthest can be attributed to the excess staples in the unpurified nanotile samples. The excess staple bands migrate a similar length and have similar intensities. The relative capacitance signatures for 1xTAE with varying Mg^2+^ are much greater than when DNA origami is introduced into the buffer. However, we observe that the relative capacitance for nanotile in 1xTAE, 20 mM Mg^2+^ is higher than nanotiles in 1xTAE, 12.5 mM Mg^2+^. A possible explanation may be that with higher salt concentrations, the nanotiles may experience salt-based adhesion and may perhaps form varying aggregates, as seen in the dim secondary gel band, which may translate as nonlinear behavior of 10 nM nanotiles in 1xTAE, 20 mM Mg^2+^. Since the interquartile range for the relative capacitance of the nanotiles annealed in 12.5 mM Mg^2+^ is the least, we infer that nanotiles in 1xTAE, 12.5 mM Mg^2+^ provide a consistent relative capacitance signature. Additionally, since our previous works^46,47^ have shown high yield for the nanotile in 1xTAE with 12.5 mM Mg^2+^, we choose 12.5 mM Mg^2+^ concentration in our 1xTAE buffers.

### DNA nanotiles concentration characterization curve shows linear increase in relative capacitance between 5 nM to 20 nM

Alongside the buffer composition, nanostructure formation is affected by the staple to scaffold ratio. In synthesis of DNA nanostructures, the scaffold is the limiting agent. The ratio of staples to scaffold, called excess ratio, is often set to be more than 1 to maximize yield of the nanostructure. The relative capacitance is noted for a solution of scaffold and staples, which are needed to make nanotiles, in 1xTAE with 12.5 mM Mg^2+^ (Figure 5(a)). It can be observed that as the excess ratio increases, so does the relative capacitance. The one-way ANOVA test supports that the relative capacitance of each excess ratio is distinct and is documented in Table S2 in Supplementary Information. It appears that an excess ratio of 5 is beneficial for capacitance studies as it provides high structure yields with minimal effects of excess ratio on the relative capacitance.

**Figure 5:**
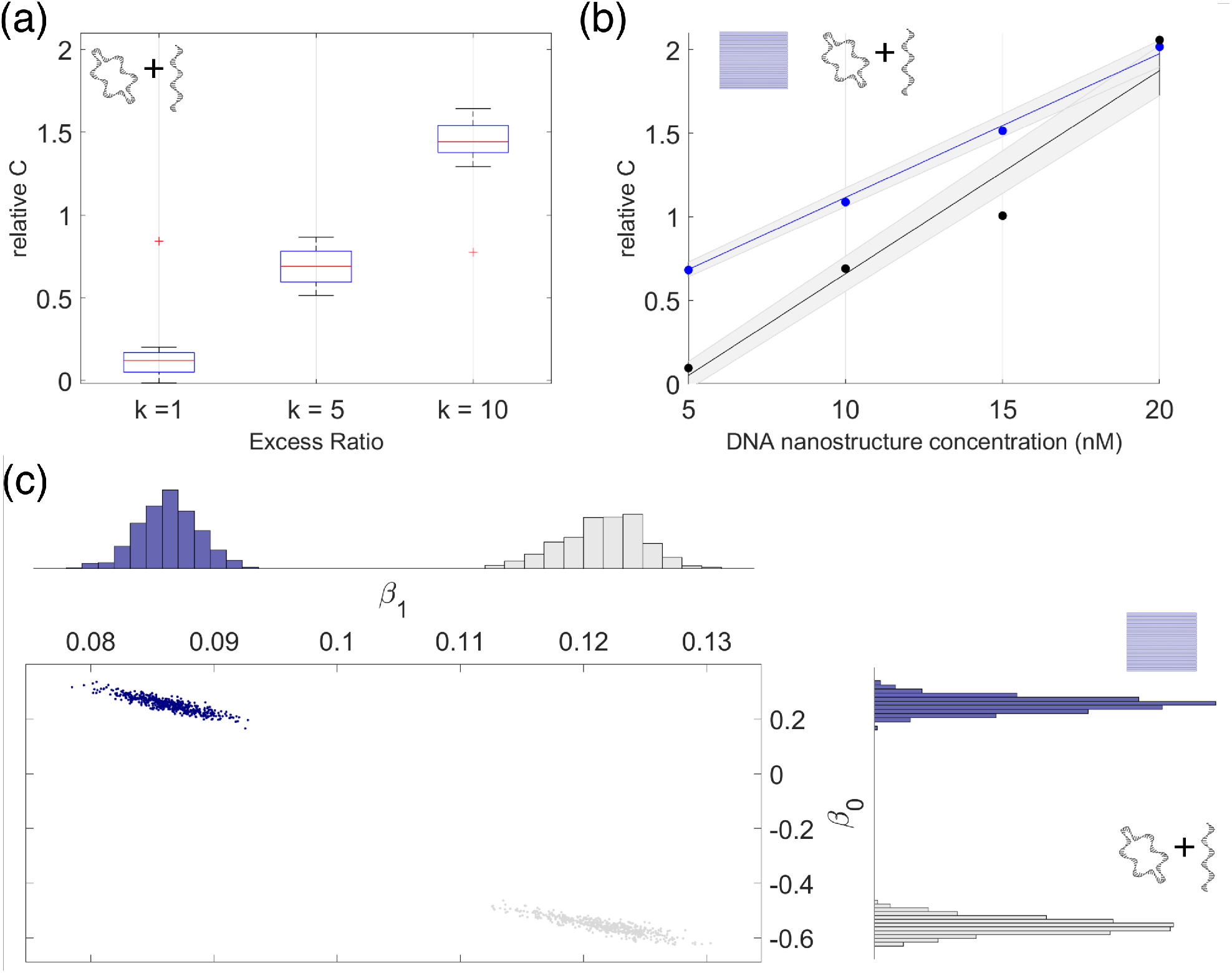
Formation assessment: relative capacitance signatures for annealed nanotiles is distinctive from pre-annealed nanotiles. (a) The relative capacitance increases with the stochiometric ratio between staples and scaffold in pre-annealed nanotiles (N=3, one-way ANOVA p=1.483e-20). (b) Concentration assessment: Nanotiles (N=3, R^2^ = 0.956; *p <* 0.01) and pre-annealed nanotiles (N=3, R^2^ = 0.924; *p <* 0.01) characteristic curves show a linear relationship between the concentration and the relative capacitance signatures. (c) Bootstrap resampling shows that characteristic curves for pre-annealed nanotiles and nanotiles are distinct.

Therefore, for the remaining studies in this work, the excess ratio is chosen to be 5. The solution of scaffold and staples needed for the nanotile with an excess ratio of 5, in 1xTAE with 12.5 mM Mg^2+^, is called the pre-annealed nanotile. The pre-annealed nanotile when subjected to a specific thermal ramp should yield well-formed nanotiles. The concentration associated with a pre-annealed nanotile is the expected concentration of nanotiles obtained when the pre-annealed solution undergoes a thermal ramp. The nanotiles are synthesized at 20 nM from the pre-annealed solution and then diluted to 5 nM, 10 nM and 15 nM of nanotiles. For each concentration, the relative capacitance of the pre-annealed nanotiles and the nanotiles marked as a median of all 16 pixels (Figure 5(b)). A linear regression is fitted to relative capacitance measured for concentrations from 5 nM to 20 nM of pre-annealed nanotiles (*r*∆*C* = *−*0.558 + 0.122*χ*; R^2^ = 0.924; *p <* 0.01), called the characteristic curve for the pre-annealed nanotiles (Figure 5(b)). Linear regression fitted to concentrations (*χ*) of unpurified nanotiles (Figure 5(b)), is called the characteristic curve for the nanotiles (*r*∆*C* = 0.254 + 0.086*χ*; R^2^ = 0.956; *p <* 0.01).

The linear fit coefficients are shown in Table 1. Using one-way ANOVA test for the pre-annealed tiles and unpurified nanotiles, the 95% confidence interval of the difference between pre-annealed and annealed nanotiles at each concentration shows no overlap for nanotile concentrations less than 15 nM as shown in Table S3 in Supplementary Information.

**Table 1:** Nanostructure characterstic curves r∆C = *β*_0_ + *β*_1_.Concentration

| Category of nanostructures | Intercept ($\beta_0$) | Slope ($\beta_1$) | Dimensions (nm) | Architecture |
| --- | --- | --- | --- | --- |
| pre-annealed nanotiles | -0.558 | 0.122 | - | - |
| nanotiles | 0.254 | 0.086 | 74 nm × 96 nm | square |
| nanospheres | 0.457 | 0.218 | 51 nm | wireframe |
| nanorods | 2.674 | 0.153 | 425 nm × 7 nm | honeycomb |

Additionally, bootstrap resampling supports that the pre-annealed nanotile and the nanotile characteristic curves are distinct. The point clouds (Figure 5(c)) presented as mean *±* S.D in Table 2, show the distribution of the slope-intercept pairs for the two characteristic curves. For the nanotile, we observe that slope (*β*_1_ = 0.086) and intercept (*β*_0_ = 0.254) of the characteristic curve are within a standard deviation of the means of the slope (*β*_1_ = 0.086*±*0.002) and intercept (*β*_0_ = 0.253*±*0.026) point clouds. For the pre-annealed nanotiles, the slope (*β*_1_ = 0.122) and intercept (*β*_0_ = *−*0.558) of the characteristic curve are within a standard deviation of the means of the slope (*β*_1_ = 0.122 *±* 0.003) and intercept (*β*_0_ = *−*0.556 *±* 0.029) point clouds.

**Table 2:** Bootstrap point-cloud parameters for slope (*β*_0_) and intercepts (*β*_1_).

| Category of nanostructures | Intercept ($\beta_0$) | Intercept ($\beta_0$) | Slope ($\beta_1$) | Slope ($\beta_1$) |
| --- | --- | --- | --- | --- |
| | linear fit | point cloud (mean ± S.D) | linear fit | point cloud (mean ± S.D) |
| pre-annealed nanotiles | -0.558 | $-0.556 \pm 0.029$ | 0.122 | $0.122 \pm 0.003$ |
| nanotiles | 0.254 | $0.255 \pm 0.028$ | 0.086 | $0.086 \pm 0.002$ |
| nanospheres | 0.457 | $0.465 \pm 0.079$ | 0.218 | $0.218 \pm 0.007$ |
| nanorods | 2.674 | $2.66 \pm 0.103$ | 0.153 | $0.153 \pm 0.008$ |

The resampling analysis and, the significant nanotile characteristic curve with a narrow confidence interval, suggests that the nanotile has a consistent relative capacitance signature. In contrast, we speculate that as the concentration of the pre-annealed nanotiles increases there is an increase in the probability of forming scaffold and staple complexes, due to non-specific binding or salt-based adhesion, causing the ‘fanned-out’ confidence interval of the pre-anneal nanotile characteristic curve. From the agarose gel electrophoresis, we are able to identify the distances the M13 scaffold and the staple strands will migrate (Figure 4(c)). The pre-annealed nanotile is a combination of scaffold and staples and so can be expected to show up as two bands in the gel - the M13 scaffold, and the excess staples farther down in the gel, as well as a possible smear of the potentially varying scaffold and staples complexes. The unpurified nanotiles also show as two bands (Figure 4(c)), the primary band close to the M13 band and the excess staples band. Thus, in agarose gel electrophoresis, it may be inconvenient to distinguish pre-annealed and post-annealed nanotiles. In contrast, relative capacitance signatures (Figure 5(b)) provide distinct characteristic curves and therefore, a clear indication of nanostructure formation.

With the relative capacitance metric, the formation and concentration of planar square-lattice DNA nanotiles is assessed based on the distinct concentration characterization curves. We consider that relative capacitance signature of a nanostructure is an combination between formation, concentration and the morphology of the nanostructure. Since the characteristic curve shows a linear correlation between concentration and the relative capacitance, we expect distinct linear characteristic curves for nanostructures with distinct morphologies.

### Distinct DNA nanostructure morphologies show distinct concentration characterization curves

Maintaining similar molecular weight to the nanotile, we investigate the relative capacitance signatures of wireframe nanospheres and honeycomb-lattice rods (Figure 2). All structures are synthesized at 20 nM, in a 1xTAE buffer with 12.5 mM of Mg^2+^ and then diluted to 5 nM, 10 nM and 15 nM concentrations using the same buffer. For each concentration, the relative capacitance for each structure is represented by the median over all the 16 pixels (Figure 6). The linear regression fits to these concentrations of unpurified structures (Figure 6(a)) are called the characteristic curve for the respective shape. Each characteristic curve is seen to be monotonic and distinct, with the linear fit coefficients as shown in Table 1. The bootstrap resampling (Figure 6(c)) shows the distribution of the slope-intercept pairs for all three characteristic curves. As noted for the nanotile, we observe that slopes and intercepts for all characteristic curves are within a standard deviation of the means of the slope and intercept point clouds, as presented in Table 2, indicating that each characteristic curve is a distinctive capacitive signature of the corresponding nanostructure.

**Figure 6:**
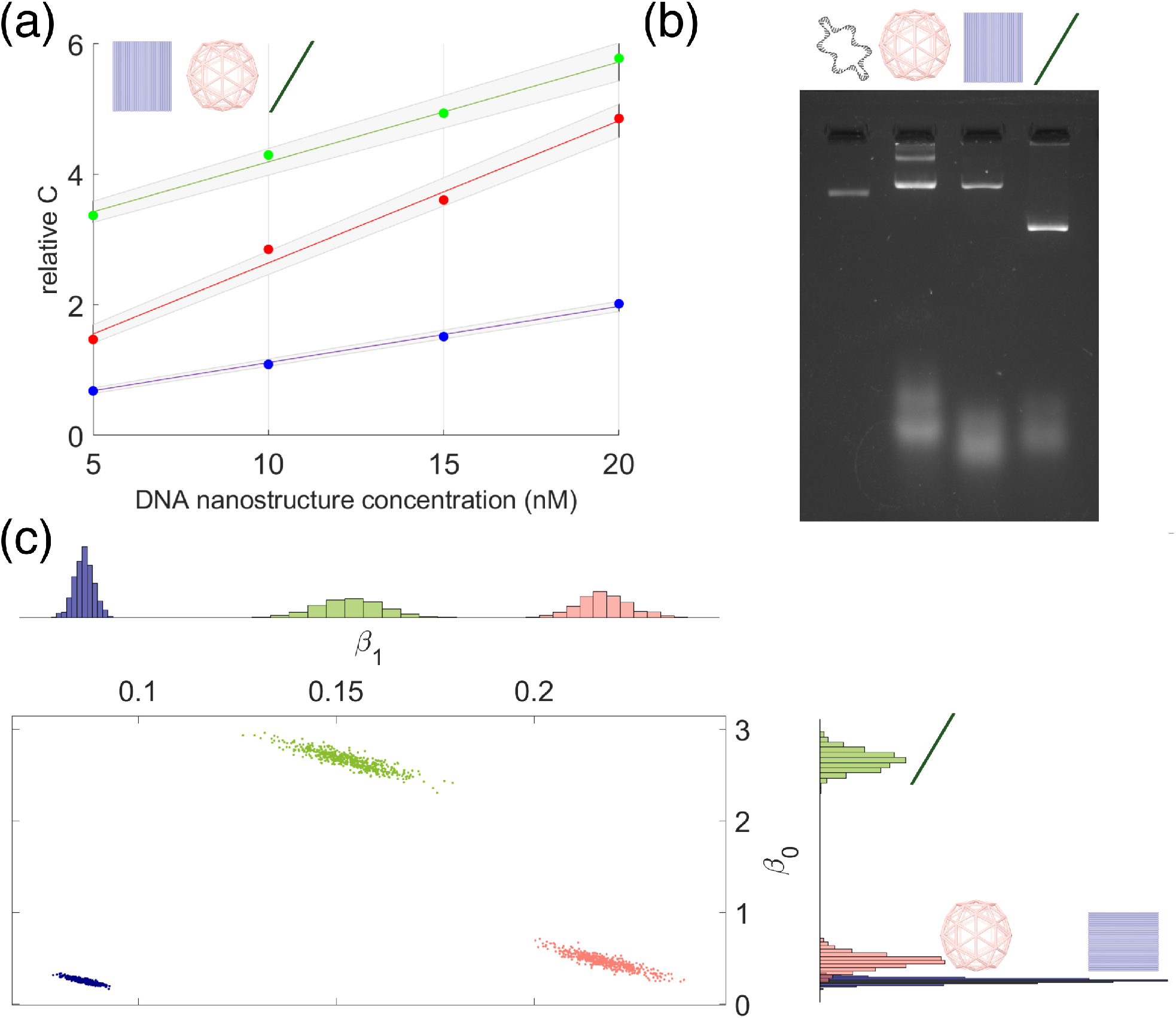
Morphology assessment: each of the three nanostructures tested have distinctive relative capacitance signatures. (a) Linear regression fits representing the characteristic curves of nanotiles (N=3, R^2^ = 0.956; *p <* 0.01), nanospheres (N=3, R^2^ = 0.931; *p <* 0.01) and nanorods (N=3, R^2^ = 0.835; *p <* 0.01). (b) Gel electrophoresis image of unpurified nanotiles, nanospheres and nanorods in 1xTAE with 12.5 mM of Mg^2+^ shows that for structures of similar molecular weight, the gel bands migrate based on geometry and dimensions of the nanostructures. (c) Bootstrap resampling shows that characteristic curves for nanotiles, nanospheres and nanorods are distinct intercepts, while the distribution of the slopes for nanospheres and nanotiles shows slight overlap.

The primary gel bands of the agarose gel electrophoresis of the nanotiles, the nanospheres and the nanorods migrate comparably while primary band of the nanorod travels the farthest (Figure 6(b)). The nanospheres have a secondary band with lighter intensity which may perhaps be an effect of nanosphere aggregation. Since the nanostructures has similar molecular weights, the gel band migration can be a result of their different morphologies. The nanotile and the nanosphere have very distinct morphologies but travel comparable distances in the gel, probably on account of their similar aspect ratios. Nevertheless, both these structures have distinctive relative capacitance signatures. Thus, relative capacitance signatures can not only provide a clear distinction between nanostructure formation and concentrations but also nanostructure morphologies.

## Conclusions

We have presented that capacitance sensing has the potential to be an affordable and rapid characterization approach to assess formation, concentration and morphology of DNA origami nanostructures. Our capacitance sensor is a 4 × 4 array of 16 electrode pairs that transduces a change in charge by changing the frequency of the output voltage. Our metric of relative capacitance has adapted our capacitance sensor to assess DNA origami nanostructures in a buffer solution of 1xTAE with MgCl_2_. We have shown that the relative capacitance is affected by the concentration of Mg^2+^ in the 1xTAE solution. In keeping with the literature,^45^ our identified operating range is 8 - 20 mM Mg^2+^ in the 1xTAE. A nonlinear change is observed in the relative capacitance as the nanotiles are synthesized in 1xTAE with 8 mM,

12.5 mM and 20 mM Mg^2+^. We interpret that Mg^2+^ in the nanostructure solution drives the change in relative capacitance. In addition, as the stoichiometric ratio of staples to scaffold in the buffer is increased, we see a monotonic increase in the relative capacitance. Therefore, maintaining the Mg^2+^ concentration at 12.5 mM and a 5-fold stoichiometric ratio we study the relative capacitance of pre-annealed and annealed nanotiles. We observe that as the concentration of pre-annealed and annealed nanotiles increases, the difference between the characteristic curves reduces and the curves almost intersect at a concentration of 20 nM.

For well-formed nanostructures, with concentrations between 5 - 20 nM, we observe distinct characteristic curves for the nanotile, nanosphere and nanorod that are designed with similar molecular weight but distinct morphology.

The viability of the capacitive characterization approach introduces a novel technology that can potentially accelerate nanostructure production. To efficiently apply this technology, our next step would be to construct a physics-based model of the capacitive response to DNA nanostructures. To explore the complete utility of this technology, a research direction is to understand how relative capacitance is affected by the interplay between nanostructure geometry, charge, architecture and even conjugations with moieties such as fluorescence dyes. Future studies can also investigate if patterning of nanostructures on the sensor surface can further impact the capacitive response. Using capacitance to characterize our DNA origami nanostructures opens potential paths for a novel technology that can complement existing characterization approaches and boost the accessibility and commercialization of DNA nanotechnology.

## Methods

### Fabrication of DNA origami nanostructures

Three nanostructures with distinct morphology (Figure 2) were generated - a nanotile, a nanorod, a nanosphere. The DNA origami nanotiles (74 × 96 nm) were constructed using a square lattice with appropriate base-pair deletions to maintain a planar 2D face as discussed in our previous work. ^46^ The DNA origami nanorods (425 nm x 7 nm) were designed using a honeycomb lattice.^48^ The DNA origami nanospheres (diameter = 51 nm) were designed as wireframe structures.^46^ These basic shape structures were synthesized by mixing the staple strands with M13mp18 scaffold (Bayou Biolabs) with 5x excess staples in 1xTAE (40 nM Tris base, 20 mM acetic acid, 1 mM EDTA). The concentration of MgCl_2_ in the 1xTAE was varied between 8 mM, 12.5 mM and 20 mM according to the study. The DNA annealing was conducted in the MiniAmp™ Plus Thermal Cycler as detailed in our previous works. ^46,48^

### Agarose Gel Electrophoresis

Unpurified DNA nanostructures, at 15 nM and 15 *µ*L, were run through 2% agarose gel for 90 minutes at 100 V, at room temperature, and visualized by adding 4 *µ*L of the SYBR Safe DNA gel stain (Invitrogen) to the gel. The gel was imaged with BioRad GelDoc G6 imaging system.

### Atomic force microscopy (AFM)

Unpurified DNA nanostructures were diluted to 1-2 nM in 1×TAE containing 12.5 mM MgCl_2_. The 10 *µ*L aliquot of nanostructures were deposited onto a freshly cleaved mica surface, incubated at room temperature in a humid chamber for 10 min. The sample was washed thrice with DI water and thoroughly blow-dried with nitrogen before imaging. The AFM was performed using the NX10 AFM system with OMCL-AC160TS tips pre-loaded onto a wafer (Park Systems Corp.), in non-contact mode (NCM). The AFM images were post-processed using the software Gwyddion (http://gwyddion.net/) and analyzed using FIJI.^49^

### Data collection

A volume of 4.5 uL of DNA nanostructures were deposited on the sensor surface in the epoxy well. For each sample deposited, data was collected for 10 minutes. The sensor surface was cleaned with DI water and dried with compressed nitrogen. The capacitance sensor response to the deposition on the sensor surface is recorded via a Teensy (https://www.pjrc.com/teensy/) microcontroller and post-processed with Python. For each deposition, the capacitance sensor responses are recorded as received from 16 independent pixels at approximately every 30 seconds, for a total of 10 minutes. The capacitance sensor responses were thresholded before statistical analysis. Multiple approaches to data processing is discussed in detail in the Supplementary Information.

### Statistical analysis

Results from 3 to 4 independent trials are shown in the figures. For each trial, DNA nanostructure samples were freshly synthesized and all samples were deposited in triplicates. Sample data were normalized using the buffer readings from the same trial and the idle sensor readings to obtain relative capacitance. Relative capacitance values were averaged over all days for each pixel to get a 16 × 1 array of relative capacitance signatures for each nanostructure category. Assuming normality over the 16 pixel array, the data for each nanostructure category was statistically analyzed with one-way ANOVA test using *anova1* function in MATLAB (version 9.12; Update 3; (R2022a). Natick, Massachusetts: The MathWorks Inc.) and compared using the Tukey-Kramer test using the *multcompare* function in MATLAB. Iteratively reweighted least squares method was used to fit a linear regression to the relative capacitance for every nanostructure concentration. Using the *fitlm* function in MATLAB, characteristic curves were generated for each nanostructure. The confidence intervals of the characteristic curves were generated from the standard errors of their respective slopes and intercepts. Parametric bootstrapping, a resampling technique, was used to determine the distinguishable nature of characteristic curves for every nanoshape category. Using the *bootstrp* function in MATLAB, 16 points were resampled from the 16 × 1 array for each nanostructure category. Regression curves were fitted to the bootstrap dataset using fitlm and represented as a scatter plot of intercept against slope. The resampling and corresponding linear regression fit was performed 500 times for each nanostructure category reported.

## Contributions

V.W., M.S.S. contributed equally as joint first authors, and M.D., R.E.T. contributed equally as joint senior authors. The manuscript was written through contributions of all authors. V.W wrote and edited the manuscript, designed and performed experimental studies, performed the data analysis; conducted literature research. M.S.S wrote the manuscript; designed and performed the experimental studies. Y.G designed the experimental studies. M.D edited the manuscript and advised the work. R.E.T wrote and edited the manuscript and advised the work.

## Data availability

All data will be made available upon request.

## Acknowledgement

The authors thank Prof. Peter Freeman, Department of Statistics, Carnegie Mellon University, Pittsburgh, PA for his insights in the statistical analysis of this data. The AFM imaging studies were performed using instrumentation funded by DURIP award FA9550-22-1-0147. This work was supported in part by CMU start-up funds and in part by the Pennsylvania Infrastructure Technology Alliance, a partnership of Carnegie Mellon, Lehigh University and the Commonwealth of Pennsylvania’s Department of Community and Economic Development (DCED).

## Supporting Information Available

The details of the nanostructure synthesis, supplementary data and derivations referenced in the paper can be found in the supplementary information on-line.

## References

(1) Dey, S.; Fan, C.; Gothelf, K. V.; Li, J.; Lin, C.; Liu, L.; Liu, N.; Nijenhuis, M. A. D.; Saccà, B.; Simmel, F. C.; Yan, H.; Zhan, P. DNA origami. 1, 1–24.

(2) Kekic, T.; Barisic, I. In silico modelling of DNA nanostructures. 18, 1191–1201.

(3) Zhang, F.; Nangreave, J.; Liu, Y.; Yan, H. Structural DNA Nanotechnology: State of the Art and Future Perspective. 136, 11198–11211.

(4) Wang, P.; Meyer, T. A.; Pan, V.; Dutta, P. K.; Ke, Y. The Beauty and Utility of DNA Origami. 2, 359–382.

(5) Strauss, M. T.; Schueder, F.; Haas, D.; Nickels, P. C.; Jungmann, R. Quantifying absolute addressability in DNA origami with molecular resolution. 9, 1600.

(6) Knappe, G. A.; Wamhoff, E.-C.; Bathe, M. Functionalizing DNA origami to investigate and interact with biological systems.

(7) Ryssy, J.; Natarajan, A. K.; Wang, J.; Lehtonen, A. J.; Nguyen, M.-K.; Klajn, R.; Kuzyk, A. Light-Responsive Dynamic DNA-Origami-Based Plasmonic Assemblies. 133, 5923–5927.

(8) DeLuca, M.; Shi, Z.; Castro, C. E.; Arya, G. Dynamic DNA nanotechnology: toward functional nanoscale devices. 5, 182–201.

(9) Ijäs, H.; Hakaste, I.; Shen, B.; Kostiainen, M. A.; Linko, V. Reconfigurable DNA Origami Nanocapsule for pH-Controlled Encapsulation and Display of Cargo. 13, 5959–5967.

(10) Pumm, A.-K.; Engelen, W.; Kopperger, E.; Isensee, J.; Vogt, M.; Kozina, V.; Kube, M.; Honemann, M. N.; Bertosin, E.; Langecker, M.; Golestanian, R.; Simmel, F. C.; Dietz, H. A DNA origami rotary ratchet motor. 607, 492–498.

(11) Wang, Y.; Le, J. V.; Crocker, K.; Darcy, M. A.; Halley, P. D.; Zhao, D.; Andrioff, N.; Croy, C.; Poirier, M. G.; Bundschuh, R.; Castro, C. E. A nanoscale DNA force spectrometer capable of applying tension and compression on biomolecules. 49, 8987–8999.

(12) Benson, E.; Carrascosa Marzo, R.; Bath, J.; Turberfield, A. J. Strategies for Constructing and Operating DNA Origami Linear Actuators. 17, 2007704.

(13) Suzuki, Y.; Kawamata, I.; Mizuno, K.; Murata, S. Large Deformation of a DNA-Origami Nanoarm Induced by the Cumulative Actuation of Tension-Adjustable Modules. 59, 6230–6234.

(14) Mills, A.; Aissaoui, N.; Maurel, D.; Elezgaray, J.; Morvan, F.; Vasseur, J. J.; Margeat, E.; Quast, R. B.; Lai Kee-Him, J.; Saint, N.; Benistant, C.; Nord, A.; Pedaci, F.; Bellot, G. A modular spring-loaded actuator for mechanical activation of membrane proteins. 13, 3182.

(15) Heuer-Jungemann, A.; Linko, V. Engineering Inorganic Materials with DNA Nanostructures. 7, 1969–1979.

(16) Herms, A.; Gänther, K.; Sperling, E.; Heerwig, A.; Kick, A.; Cichos, F.; Mertig, M. Concept, synthesis, and structural characterization of DNA origami based self-thermophoretic nanoswimmers. 214, 1600957.

(17) Wijesekara, P.; Liu, Y.; Wang, W.; Johnston, E. K.; Sullivan, M. L. G.; Taylor, R. E.; Ren, X. Accessing and Assessing the Cell-Surface Glycocalyx Using DNA Origami. 21, 4765–4773.

(18) Kosuri, P.; Altheimer, B. D.; Dai, M.; Yin, P.; Zhuang, X. Rotation tracking of genome-processing enzymes using DNA origami rotors. 572, 136–140.

(19) Bian, X.; Zhang, Z.; Xiong, Q.; De Camilli, P.; Lin, C. A programmable DNA-origami platform for studying lipid transfer between bilayers. 15, 830–837.

(20) Weiden, J.; Bastings, M. M. DNA origami nanostructures for controlled therapeutic drug delivery. 52, 101411.

(21) Shen, L.; Wang, P.; Ke, Y. DNA Nanotechnology-Based Biosensors and Therapeutics. 10, 2002205.

(22) Castro, C. E.; Kilchherr, F.; Kim, D.-N.; Shiao, E. L.; Wauer, T.; Wortmann, P.; Bathe, M.; Dietz, H. A primer to scaffolded DNA origami. 8, 221–229.

(23) Jun, H.; Wang, X.; Parsons, M.; Bricker, W.; John, T.; Li, S.; Jackson, S.; Chiu, W.; Bathe, M. Rapid prototyping of arbitrary 2D and 3D wireframe DNA origami. 49, 10265–10274.

(24) Wagenbauer, K. F.; Engelhardt, F. A. S.; Stahl, E.; Hechtl, V. K.; Stömmer, P.; Seebacher, F.; Meregalli, L.; Ketterer, P.; Gerling, T.; Dietz, H. How We Make DNA Origami. 18, 1873–1885.

(25) Mathur, D.; Medintz, I. L. Analyzing DNA Nanotechnology: A Call to Arms For The Analytical Chemistry Community. 89, 2646–2663.

(26) Wang, J.; Wei, Y.; Zhang, P.; Wang, Y.; Xia, Q.; Liu, X.; Luo, S.; Shi, J.; Hu, J.; Fan, C.; Li, B.; Wang, L.; Zhou, X.; Li, J. Probing Heterogeneous Folding Pathways of DNA Origami Self-Assembly at the Molecular Level with Atomic Force Microscopy. 22, 7173–7179.

(27) Lee Tin Wah, J.; David, C.; Rudiuk, S.; Baigl, D.; Estevez-Torres, A. Observing and Controlling the Folding Pathway of DNA Origami at the Nanoscale. 10, 1978–1987.

(28) Rothemund, P. W. K. Folding DNA to create nanoscale shapes and patterns. 440, 297–302.

(29) Marrs, J.; Lu, Q.; Pan, V.; Ke, Y.; Hihath, J. Structure-Dependent Electrical Conductance of DNA Origami Nanowires. n/a, e202200454.

(30) Kabiri, Y.; Ravelli, R. B. G.; Lehnert, T.; Qi, H.; Katan, A. J.; Roest, N.; Kaiser, U.; Dekker, C.; Peters, P. J.; Zandbergen, H. Visualization of unstained DNA nanostructures with advanced in-focus phase contrast TEM techniques. 9, 7218.

(31) Dietz, H.; Douglas, S. M.; Shih, W. M. Folding DNA into Twisted and Curved Nanoscale Shapes. 325, 725–730.

(32) Fu, D.; Pradeep Narayanan, R.; Prasad, A.; Zhang, F.; Williams, D.; Schreck, J. S.; Yan, H.; Reif, J. Automated design of 3D DNA origami with non-rasterized 2D curvature. 8, eade4455.

(33) Bai, X.-c.; Martin, T. G.; Scheres, S. H. W.; Dietz, H. Cryo-EM structure of a 3D DNA-origami object. 109, 20012–20017.

(34) Kube, M.; Kohler, F.; Feigl, E.; Nagel-Yüksel, B.; Willner, E. M.; Funke, J. J.; Gerling, T.; Stömmer, P.; Honemann, M. N.; Martin, T. G.; Scheres, S. H. W.; Dietz, H. Revealing the structures of megadalton-scale DNA complexes with nucleotide resolution. 11, 6229.

(35) Schnitzbauer, J.; Strauss, M. T.; Schlichthaerle, T.; Schueder, F.; Jungmann, R. Superresolution microscopy with DNA-PAINT. 12, 1198–1228.

(36) Hartl, C.; Frank, K.; Amenitsch, H.; Fischer, S.; Liedl, T.; Nickel, B. Position Accuracy of Gold Nanoparticles on DNA Origami Structures Studied with Small-Angle X-ray Scattering. 18, 2609–2615.

(37) Bruetzel, L. K.; Walker, P. U.; Gerling, T.; Dietz, H.; Lipfert, J. Time-Resolved Small-Angle X-ray Scattering Reveals Millisecond Transitions of a DNA Origami Switch. 18, 2672–2676.

(38) Baker, M. A. B.; Tuckwell, A. J.; Berengut, J. F.; Bath, J.; Benn, F.; Duff, A. P.; Whitten, A. E.; Dunn, K. E.; Hynson, R. M.; Turberfield, A. J.; Lee, L. K. Dimensions and Global Twist of Single-Layer DNA Origami Measured by Small-Angle X-ray Scattering. 12, 5791–5799.

(39) Wang, X.; Mao, Z.; Chen, R.; Li, S.; Ren, S.; Liang, J.; Gao, Z. Self-assembled DNA origami-based duplexed aptasensors combined with centrifugal filters for efficient and rechargeable ATP detection. 211, 114336.

(40) Hanke, M.; Grundmeier, G.; Keller, A. Direct visualization of the drug loading of single DNA origami nanostructures by AFM-IR nanospectroscopy. 14, 11552–11560.

(41) Iäas, H.; Liedl, T.; Linko, V.; Posnjak, G. A label-free light-scattering method to resolve assembly and disassembly of DNA nanostructures. 121, 4800–4809.

(42) Prakash, S. B.; Abshire, P. On-Chip Capacitance Sensing for Cell Monitoring Applications. IEEE Sensors Journal 2007, 7, 440–447.

(43) Roodhuizen, J. A. L.; Hendrikx, P. J. T. M.; Hilbers, P. A. J.; de Greef, T. F. A.; Markvoort, A. J. Counterion-Dependent Mechanisms of DNA Origami Nanostructure Stabilization Revealed by Atomistic Molecular Simulation. 13, 10798–10809.

(44) Senevirathna, B. P.; Lu, S.; Dandin, M. P.; Basile, J.; Smela, E.; Abshire, P. A. Real-Time Measurements of Cell Proliferation Using a Lab-on-CMOS Capacitance Sensor Array. 12, 510–520.

(45) Kielar, C.; Xin, Y.; Shen, B.; Kostiainen, M. A.; Grundmeier, G.; Linko, V.; Keller, A. On the Stability of DNA Origami Nanostructures in Low-Magnesium Buffers. 57, 9470–9474.

(46) Liu, Y.; Wijesekara, P.; Kumar, S.; Wang, W.; Ren, X.; Taylor, R. E. The effects of overhang placement and multivalency on cell labeling by DNA origami. 13, 6819–6828.

(47) Roka-Moiia, Y.; Walawalkar, V.; Liu, Y.; Italiano, J. E.; Slepian, M. J.; Taylor, R. E. DNA Origami–Platelet Adducts: Nanoconstruct Binding without Platelet Activation. 33, 1295–1310.

(48) Wang, W.; Hayes, P. R.; Ren, X.; Taylor, R. E. Synthetic cell armor made of DNA origami. (Submitted).

(49) Schindelin, J. et al. Fiji: an open-source platform for biological-image analysis. 9, 676–682.

