## Supplementary Information for "Capacitance measurements for assessing DNA origami nanostructures"

§*equal contribution*

All nanostructure caDNAno files are provided in the Supplementary Information of our previous works.[1,2]

| name (5'-3') | Sequence |
|---|---|
| seam01 | TCAGGAGGTTTAGTACCGCCACCCTCAGAAC |
| seam02 | TTTGCTAAGTAAATGAATTTTCTGAGTGCCTT |
| seam03 | GAGTAACAGTTTTAACGGGGTCTATGGGAT |
| seam04 | TAACCGATATGACAACAACCATCGCACCACCC |
| seam05 | TCAGAGCCGCCACCCTCAGAGCCCCACGCA |
| seam06 | CACTAAAATAAAACGAAAGAGGCAACGTCACC |
| seam07 | AATGAAATTAGCAAGGCCGGAAAAAGAATA |
| seam08 | TGACCTTCTACAGACCAGGCGCATACCACGGA |
| seam09 | ATAAGTTAAGAAACGCAAAGACAGGCTGGC |
| seam10 | AAAATCTAATACCAGTCAGGACGTGCAAGAAA |
| seam11 | CAATGAAAGCCCAATAATAAGATGGGAAGA |
| seam12 | GGGGTAATTTTGCAAAAGAAGTTTAAATAAAC |
| seam13 | AGCCATAAATTTGCCAGTTACATGCCAGAG |
| seam14 | TTCGAGCTAAGACTTCAAATATCGATCGTAGG |
| seam15 | AATCATTGCCGTTTTTATTTTCCGTTTTAA |
| seam16 | AGTAGATTAGTTGATTCCCAATTCTTCTGTCC |
| seam17 | AGACGACCAAAAGGTAAAGTAATGCGAACG |
| seam18 | CCCTGTAATCGGTTGTACCAAAAATAAGGCGT |
| seam19 | TAAATAACGACCGTGTGATAAACATTATGA |
| seam20 | AGGCTATCTAGCTATTTTTGAGAGAAGACGCT |
| seam21 | GAGAAGAGCGATAGCTTAGATTATCTACAA |
| seam22 | ACGCCATCCAGCTCATTTTTTAACAGGCGAAT |
| seam23 | TATTCATTACAAAATCGCGCAGCAATAGGA |
| seam24 | AGCTTTCCCCTCAGGAAGATCGCACATCAATA |
| seam25 | TAATCCTCAGATGATGGCAATTCTCCAGCC |
| seam26 | GGTCGACTCAGTGCCAAGCTTGCAGGAGCACT |
| seam27 | AACAACTTCTAAAATATCTTTATGCCTGCA |
| seam28 | TTAATGAAGGGAAACCTGTCGTGCATTAAAAA |
| seam29 | TACCGAACCCTAAAACATCGCCCAGCTGCA |
| seam30 | CCCGAGATATCCCTTATAAATCAAAAACGCTC |
| seam31 | ATGGAAACCATTGCAACAGGAAAAGAATAG |
| seam32 | GAGCTTGACGGGGAAAAAAGGGATTTTAGAC |
| X01Y01 | TAGGAACCCATGTACACAACGCC |
| X01Y02 | TGTAGCATAATTTTTTCACGTTCCTTTAAT |
| X01Y03 | TGTATCGAGCGAAAGACAGCATTGAGGACT |
| X01Y04 | AAAGACTCATCGCCTGATAAATTTAGCCGG |
| X01Y05 | AACGAGGATTCAGTGAATAAGGTAAATTGG |
| X01Y06 | GCTTGAGTAGGAATACCACATTTTACGAGG |
| X01Y07 | CATAGTACCCCTCAAATGCTTTCAAAAATC |
| X01Y08 | AGGTCTTTCCTTTTGATAAGAGCTGAATAT |
| X01Y09 | AATGCTGGGGCGCGAGCTGAAATTAACATC |
| X01Y10 | CAATAAATTTAAATGCAATGCCGAGAAAGG |
| X01Y11 | CCGGAGAATGTCAATCATATGTCAGGAAGA |
| X01Y12 | TTGTATAACAACCCGTCGGATTTGGGATAG |
| X01Y13 | GTCACGTTGGGAAGGGCGATCGGAAAGGGG |
| X01Y14 | GATGTGCAATTGTTATCCGCTCAAAGTGTA |
| X01Y15 | AAGCCTGAGTGAGACGGGCAACGAGTTGCA |
| X01Y16 | GCAAGCGCAAAGGGCGAAAAACCATCACCCAAATCAA |
| X02Y01 | GCGAATAATTCCACAGACAGCCCTGGGATAGCAAGCCCAA |
| X02Y02 | CCCTCAGCGTTTATCAGCTTGCTTAAGGAATT |
| X02Y03 | ATTTGTATTTTTCATGAGGAAGTTGATCGTCA |
| X02Y04 | AGCTGCTCCGCAGACGGTCAATCACAACGGAG |
| X02Y05 | TTGAGATTATGGTTTAATTTCAACCGTAACAA |
| X02Y06 | ATTGAATCAGAGCAACACTATCATTTCATCAG |
| X02Y07 | TTAATTGCTACCCTGACTATTATAAAATATTC |
| X02Y08 | TTCATTTGTAGCTCAACATGTTTTGAGTACCT |
| X02Y09 | TCATATATTCATACAGGCAAGGCAGCTATATT |
| X02Y10 | AAACTAGCCAGTCAAATCACCATCTAGAACCC |
| X02Y11 | AGCGAGTAAGCAAATATTTAAATTTAATCGTA |
| X02Y12 | GCAACTGTTGGTGTAGATGGGCGCTAAATGTG |
| X02Y13 | CTGTGTGATGCAAGGCGATTAAGTCAGGCTGC |
| X02Y14 | TTTTCACCGGGTGCCTAATGAGTGGCTGTTTC |
| X02Y15 | TCCAACGTGTCCACGCTGGTTTGCGGTTTTTC |
| X02Y16 | GTTTTTTGGGGTCGAGACGTGGAC |
| X03Y01 | ACCACCCTCATTTTCACATAGTTAGCGTAACGTAGAAAGG |
| X03Y02 | AACAACTATCGAGGTGAATTTCTTAAGGCCGC |
| X03Y03 | TTTTGCGGTCCATTAAACGGGTAAAGCGCGAA |
| X03Y04 | ACAAAGTATAAGGGAACCGAACTGTCATTACC |
| X03Y05 | CAAATCAATTTAATCATTGTGAATATTACAGG |
| X03Y06 | TAGAAAGAAACCCTCGTTTACCAGACTGCGGA |
| X03Y07 | ATCGTCATGTCAGAAGCAAAGCGGACAGGTCA |
| X03Y08 | GGATTAGAAAATATGCAACTAAAGGTCAATAA |
| X03Y09 | CCTGTTTAAAGAATTAGCAAAATTCAAGGATA |
| X03Y10 | AAAATTTTAATATGATATTCAACCGAGAATCG |
| X03Y11 | ATGAACGGGTAAACGTTAATATTTCAGCTTTC |
| X03Y12 | ATCAACATATCGTAACCGTGCATCAGCGCCAT |
| X03Y13 | TCGCCATTTGGGTAACGCCAGGGTCGTAATCA |
| X03Y14 | TGGTCATAAGCTAACTCACATTAATATTGGGC |
| X03Y15 | GCCAGGGTCCCAGCAGGCGAAAATAGTCCACT |
| X03Y16 | ATTAAAGAGTGCCGTAAAGCACTA |
| X04Y01 | AGTGAGAAATCTAAAGTTTTGTCGCCACCCTCAGAGCC |
| X04Y02 | GGGAGTTAAAACAGCTTGATACCTCAGCGG |
| X04Y03 | TTATACCAAATACGTAATGCCACGCTTGCA |
| X04Y04 | CCGGATATACCAACTTTGAAAGACCAGCGA |
| X04Y05 | ACAACATTTACCTTATGCGATTTACAAGAA |
| X04Y06 | CGTCCAATACGACGATAAAAACCTAACGGA |
| X04Y07 | AAACTCCAATTGCATCAAAAAGATGGATAG |
| X04Y08 | CGCAAATGTACGGTGTCTGGAAGCGGAAGC |
| X04Y09 | TTTCAACGAAGCAATAAAGCCTCTACATTT |
| X04Y10 | GCAAACAAGTTCTAGCTGATAAAGCCTTTA |
| X04Y11 | CCTGTAGCTGTTAAAATTCGCATGTCTGGA |
| X04Y12 | CCAGGCAATGCCAGTTTGAGGGGTGGCCTT |
| X04Y13 | CTCGAATTTTTCCCAGTCACGACCCGGAAA |
| X04Y14 | GGTTTGCGTTGCGTTGCGCTCACTACCGAG |
| X04Y15 | GGAACAAGCCTGTTTGATGGTGGGAGAGGC |
| X04Y16 | AATCGGAACCCTAAACCAGTTT |
| X05Y01 | CGCCACCCTCAGAACCGTCTTTCCAGACGTTAACAACTTT |
| X05Y02 | CAACAGTTGATAGTTGCGCCGACAATATTCGG |
| X05Y03 | TCGCTGAGTACGAAGGCACCAACCCACTCATC |
| X05Y04 | TTTGACCCGGACAGATGAACGGTGATCAAGAG |
| X05Y05 | TAATCTTGTAAGAACTGGCTCATTCGTTAATA |
| X05Y06 | AAACGAACAAAATAGCGAGAGGCTAGTAAAAT |
| X05Y07 | GTTTAGACTTAAGAGGAAGCCCGATCAAAGCG |
| X05Y08 | AACCAGACTTTCATTCCATATAACTAGTTTGA |
| X05Y09 | CCATTAGAAGAGCATAAAGCTAAATACTTTTG |
| X05Y10 | CGGGAGAATTAATGCCGGAGAGGGAGGTCATT |
| X05Y11 | GCCTGAGATAAATTTTTGTTAAATAAAAATAA |
| X05Y12 | TTCGCGTCACGACGACAGTATCGGGGCACCGC |
| X05Y13 | TTCTGGTGGTTGTAAAACGACGGCCTAGAGGA |
| X05Y14 | TCCCCGGGTGCCCGCTTTCCAGTCTCGGCCAA |
| X05Y15 | CGCGCGGGTTCCGAAATCGGCAAAAGGGTTGA |
| X05Y16 | GTGTTGTTGGGAGCCCCCGATTTA |
| X06Y01 | TGGTAATAAGTGCCCGTATAAACAAGGTGTATCACCGTAC |
| X06Y02 | CTCAGAACCGCCACCAGAACCACCGAGTGTAC |
| X06Y03 | CATTACCACCATCGATAGCAGCACCCGCCACC |
| X06Y04 | ACATATAATATTTTGTCACAATCAAGTAGCAC |
| X06Y05 | TTGAGTTAATAGCAATAGCTATCTAGGTGGCA |
| X06Y06 | CAGAGCCTTTATTTATCCCAATCCCACAAGAA |
| X06Y07 | ACAAGCAAACCGCGCCCAATAGCACGTCTTTC |
| X06Y08 | AGTACCGAGACAATAAACAACATGCATCGAGA |
| X06Y09 | TGAAATACGAATAAACACCGGAATGAATATAA |
| X06Y10 | AAAACATAGTCAATAGTGAATTTATAATGGTT |
| X06Y11 | TACCAAGTTTCAATTACCTGAGCAAATCCTTG |
| X06Y12 | CTGATTATGATTGTTTGGATTATAGCTTTGAA |
| X06Y13 | GAAGGTTAAATAGATTAGAGCCGTTCATATTC |
| X06Y14 | ACTGATAGCGAACCACCAGCAGAAGAATTGAG |
| X06Y15 | ACCGCCAGTACCTACATTTTGACGATGCGCGA |
| X06Y16 | AGGAACGGTACGCCAGACAATATT |
| X07Y01 | AAGTATAGCCCGGAATGTTAATGCCCCCTGCCGCTTTTGA |
| X07Y02 | TGATACAGACCAGAGCCGCCGCCAACCGCCTC |
| X07Y03 | CCTCAGAGCGTAATCAGTAGCGACAGCCAGCA |
| X07Y04 | AAATCACCATAGAAAATTCATATGGAAAATAC |
| X07Y05 | ATACATAATACCGAAGCCCTTTTTTATCAGAG |
| X07Y06 | AGATAACCAAATAAGAAACGATTTTACCAACG |
| X07Y07 | CTAACGAGAGCAAATCAGATATAGAACCAAGT |
| X07Y08 | ACCGCACTTTCAGCTAATGCAGAACGAGCCAG |
| X07Y09 | TAATAAGACATAATTACTAGAAAATCTTCTGA |
| X07Y10 | CCTAAATTTCAAAATCATAGGTCTATTAATTT |
| X07Y11 | TCCCTTAGAAAGAAGATGATGAAAACGGATTC |
| X07Y12 | GCCTGATTCTTCTGAATAATGGAAAGGAGCGG |
| X07Y13 | AATTATCACAATAGATAATACATTAATCAACA |
| X07Y14 | GTTGAAAGGATAAAACAGAGGTGATGGCTATT |
| X07Y15 | AGTCTTTACTCAATCGTCTGAAATGCTGGTAA |
| X07Y16 | TATCCAGAAATCCTGAGAAGTGTT |
| X08Y01 | CATACATGTATTTCGGAACCTATCGAGAGGGTTGATAT |
| X08Y02 | CCACCGGAGCATTGACAGGAGGTTAAGCGT |
| X08Y03 | GGAATTAGAGAATCAAGTTTGCCAGAGCCA |
| X08Y04 | CAAACGTAGTTTACCAGCGCCAACCATTTG |
| X08Y05 | AGCGCTAAAAGAAAAGTAAGCAGATGTTAG |
| X08Y06 | CTGAATCTTTTGTTTAACGTCAAGTAATTG |
| X08Y07 | GGGTATTAAAGGCTTATCCGGTATTTTATC |
| X08Y08 | GGCATTTTCGCGCCTGTTTATCACAAGAAC |
| X08Y09 | TAATTTCAAGCCTGTTTAGTATCAGGCAGA |
| X08Y10 | CGCTATTAGAGAGACTACCTTTTTTTTAGT |
| X08Y11 | AAACAATACAAACATCAAGAAAAAAAATCGT |
| X08Y12 | CCACCAGAGGGTTAGAACCTACCTCGGGAG |
| X08Y13 | AGTTGGCATGAGGATTTAGAAGTAAAGAAA |
| X08Y14 | TTTTTGAAGGCGGTCAGTATTAATCTGGTC |
| X08Y15 | TCGGCCTTGGATTATTTACATTGGACAATA |
| X08Y16 | TTTATAATCAGTGAGCAAACTA |
| X09Y01 | GGCGGATAAGTGCCGTTATTCTGAAACATGAAGAATTTAC |
| X09Y02 | CGTTCCAGTGAGGCAGGTCAGACGCAAAATCA |
| X09Y03 | CCGGAACCTTTAGCGTCAGACTGTCCGTCACC |
| X09Y04 | GACTTGAGAGACAAAAGGGCGACACTCCTTAT |
| X09Y05 | TACGCAGTATAGCCGAACAAAGTTTGAACAAA |
| X09Y06 | GTCAGAGGAAATGAAAATAGCAGCTTGCACCC |
| X09Y07 | AGCTACAATTCTAAGAACGCGAGGTCTTTCCT |
| X09Y08 | TATCATTCACAATAGATAAGTCCTCGCCAACA |
| X09Y09 | TGTAATTTATATGCGTTATACAAAAACTTTTT |
| X09Y10 | CAAATATATAACCTCCGGCTTAGGATAACCTT |
| X09Y11 | GCTTCTGTCAAAATTAATTACATTAACAGTAC |
| X09Y12 | CTTTTACAATATCAAAATTATTTGTATCATTT |
| X09Y13 | TGCGGAACATTAGACTTTACAAACAAACCCTC |
| X09Y14 | AATCAATACACCGCCTGCAACAGTAAGAATAC |
| X09Y15 | GTGGCACAGCAGATTCACCAGTCACCTGAGTA |
| X09Y16 | GAAGAACTGCCACCGAGTAAAAGA |
| X10Y01 | CAGTCTCTAGTATTAAGAGGCTGAGTTTTGCTCAGTACCA |
| X10Y02 | TTCATAATATTGGCCTTGATATTCGGAAAGCG |
| X10Y03 | AATTATCAAGCGCGTTTTCATCGGGCCATCTT |
| X10Y04 | GATTAAGATTCAACCGATTGAGGGTAAAGGTG |
| X10Y05 | GAACACCCACCAGAAGGAAACCGAACTGGCAT |
| X10Y06 | TTGCTATTCTTTACAGAGAGAATAAATTAACT |
| X10Y07 | ATCGGCTGCGTTTTAGCGAACCTCAAGATTAG |
| X10Y08 | TTTAACAAGAACAAGAAAAATAATAATCAATA |
| X10Y09 | CGCGAGAATTCTTACCAGTATAAATCGCCATA |
| X10Y10 | GTGAGTGATTGGGTTATATAACTAACAAAGAA |
| X10Y11 | TATACAGTTAACAATTTCATTTGACAATATAT |
| X10Y12 | AGTAACATCACGTAAAACAGAAATCAGATGAA |
| X10Y13 | CAAATATCAATTCGACAACTCGTAAAAGTTTG |
| X10Y14 | GAAAGCGTGCCACGCTGAGAGCCACTGAACCT |
| X10Y15 | ATCACTTGCACGACCAGTAATAAATCTGACCT |
| X10Y16 | GTCTGTCCATCACGCAGTAATAAC |

Figure S1: Staples sequences for synthesis of nanotile.

| name(5'-3') | sequence |
|---|---|
| helix1-helix14 | CCACCCTCAGAGCCACCACCCTCATAGCTATCTTACCGAAGCCCT |
| helix2-helix36 | AGCCACCACCGGAACCGCCTCCCACTATATGTAAATGCTGATGCAAATCC |
| helix3-helix11 | TTATTAGCGTTTGCCATCTTTTCATGTTAGCAAACGTAGAAAAT |
| helix4-helix32 | CGTCAGACTGTAGCGCGTTTTCAATCATATGCGTTATACAAATTCTTACC |
| helix5-helix7 | GATAGCAGCACCGTAATCAGTAGAAAGGTGAATTATCACCGTCACCG |
| helix6-helix28 | ACCATTACCATTAGCAAGGCCATGTTCAGCTAATGCAGAACGCGC |
| helix7-helix24 | ACTTGAGCCATTTGGGAATTAGATTTCATCGTAGGAATCATTAC |
| helix8-helix10 | AATATTGACGGAAATTATTCATTTATAAAAGAAACGCAAAGACACCA |
| helix9-helix20 | AGCGCCAAAGACAAAAGGGCGACATTCGAGCGTCTTTCCAGAGCCTAATT |
| helix10-helix12 | CGGAATAAGTTTATTTTGTCACAGGAATACCCAAAAGAACTGGCATG |
| helix11-helix4 | ACATACATAAAGGTGGCAACACGACAGAATCAAGTTTGCCTTTAG |
| helix12-helix13 | ATTAAGACTCCTTATTACGCAGTAGCCGAACAAAGTTACCAGAAGGA |
| helix13-helix15 | AACCGAGGAAACGCAATAATAACAATTGAGTTAAGCCCAATAATAAG |
| helix14-helix2 | TTTTAAGAAAAGTAAGCAGATATAATCAAAATCACCGGAACCAG |
| helix15-helix129 | AGCAAGAAACAATGAAATAGCAATACCGTTCCAGTAAGCGTCATACATGG |
| helix16-helix18 | ATATCAGAGAGATAACCCACAAGCAAAAATGAAAATAGCAGCCTTTA |
| helix17-helix125 | ATTAACTGAACACCCTGAACAGATTAGCGGGGTTTTGCTCAGTAC |
| helix18-helix19 | CAGAGAGAATAACATAAAAACAGATATTATTTATCCCAATCCAAATA |
| helix19-helix9 | AGAAACGATTTTTTGTTTAACGTATCAATAGAAAATTCATATGGTTTACC |
| helix20-helix123 | TGCCAGTTACAAAATAAACAGCCAGGTTTAGTACCGCCACCCTC |
| helix21-helix23 | ATCCTGAATCTTACCAACGCTAACAGATATAGAAGGCTTATCCGGTA |
| helix22-helix119 | TGAAGCCTTAAATCAAGATTAGTTGCAGTTTTGTCGTCTTTCCAGACGTT |
| helix23-helix25 | TTCTAAGAACGCGAGGCGTTTTAATTAAACCAAGTACCGCACTCATC |
| helix24-helix8 | CGCGCCCAATAGCAAGCAAATCAACCGATTGAGGGAGGGAAGGTA |
| helix25-helix26 | GAGAACAAGCAAGCCGTTTTTATAACCAATCAATAATCGGCTGTCTT |
| helix26-helix114 | TCCTTATCATTCCAAGAACGGGTATAGTTGCGCCGACAATGACAACAACC |
| helix27-helix6 | ATCCTAATTTACGAGCATGTAGAGCCAGCAAAATCACCAGTAGC |
| helix28-helix110 | CTGTTTATCAACAATAGATAAGTCCATTAAACGGGTAAAATACGTAATGC |
| helix29-helix31 | TCCAGACGACGACAATAAACAACGGCTTAATTGAGAATCGCCATATT |
| helix30-helix80 | GTAATAAGAGAATATAAAGTAACCTGTCGTGCCAGCTGCATT |
| helix31-helix33 | TAACAACGCCAACATGTAATTTATAAGAATAAACACCGGAATCATAA |
| helix32-helix5 | AGTATAAAGCCAACGCTCAACAGTAGGGAAACGTCACCAATGAAACCATC |
| helix33-helix35 | TTACTAGAAAAAGCCTGTTTAGTACTTTTTCAAATATATTTTAGTTA |
| helix34-helix79 | TGTGATAAATAAGGCGTTAAAGGAGAGGCGGTTTGCGTATTGGGC |
| helix35-helix37 | ATTTCATCTTCTGACCTAAATTTGTCTGAGAGACTACCTTTTTAACC |
| helix36-helix3 | AATCGCAAGACAAAGAACGCGAGAAATCGGCATTTTCGGTCATAGCCCCC |
| helix37-helix39 | TCCGGCTTAGGTTGGGTTATATATTAATTTTCCCTTAGAATCCTTGA |
| helix38-helix77 | GTGAATTTATCAAAATCATAGAGAGTTGCAGCAAGCGGTCCACGC |
| helix39-helix41 | AAACATAGCGATAGCTTAGATTACATTTAACAATTTCATTTGAATTA |
| helix40-helix1 | TTGCTTCTGTAAATCGTCGCTATTAATCAGAGCCGCCACCCTCAGAACCG |
| helix41-helix42 | CCTTTTTTAATGGAAACAGTACAAGCAAAAGAAGATGATGAAACAAA |
| helix42-helix75 | CATCAAGAAAACAAAATTAATTAAAAGAATAGCCCGAGATAGGG |
| helix43-helix45 | GAATTATTCATTTCAATTACCTGATTGCGTAGATTTTCAGGTTTAAC |
| helix44-helix71 | CGGATTCGCCTGATTGCTTTGAATACGGGAGCCCCCGATTTAGAGCTTGA |
| helix45-helix47 | GTCAGATGAATATACAGTAACAGTCCTGATTGTTTGGATTATACTTC |
| helix46-helix131 | ACGTAAAACAGAAATAAAGAAAGGTTGAGGCAGGTCAGACGA |
| helix47-helix127 | TGAATAATGGAAGGGTTAGAACCAACAGTTAATGCCCCCTGCCT |
| helix48-helix69 | ATGATGGCAATTCATCAATATAACAAGTGTAGCGGTCACGCTGCGCGTAA |
| helix49-helix51 | GAAACCACCAGAAGGAGCGGAATGAGGATTTAGAAGTATTAGACTTT |
| helix50-helix57 | ATTAATTTTAAAAGTTTGAGTTGATAGCCCTAAAACATCGCC |
| helix51-helix53 | ACAAACAATTCGACAACTCGTATTGGCAAATCAACAGTTGAAAGGAA |
| helix52-helix126 | TAGAGCCGTCAATAGATAATACATTTAAACATGAAAGTATTAAGAGGCTG |
| helix53-helix121 | TTGAGGAAGGTTATCTAAAATATTAGGAACCCATGTACCGTAACACTGAG |
| helix54-helix56 | CCCTCAATCAATATCTGGTCAGTACCAGCAGAAGATAAAACAGAGGT |
| helix55-helix100 | CAGCAAATGAAAAATCTAAAGCATCATGGCTCATTATACCAGTCAGGACG |
| helix56-helix58 | GAGGCGGTCAGTATTAACACCGCGGCACAGACAATATTTTTGAATGG |
| helix57-helix50 | ATTAAAAATACCGAACGAACCTAAATCCTTTGCCCGAACGTT |
| helix58-helix66 | CTATTAGTCTTTAATGCGCGAACAAAGGGATTTTAGACAGGAACGGTACG |
| helix59-helix98 | GACCTGAAAGCGTAAGAATACGTAGTTGAGATTTAGGAATACCA |
| helix60-helix62 | ACCAGTAATAAAAGGGACATTCTACAATATTACCGCCAGCCATTGCA |
| helix61-helix94 | CGTCTGAAATGGATTATTTACGCGTCCAATACTGCGGAATCG |
| helix62-helix88 | ACAGGAAAAACGCTCATGGAAATTTGCGGATGGCTTAGAGCTTAATTGCT |
| helix63-helix65 | CGGCCTTGCTGGTAATATCCAGAATCAGTGAGGCCACCGAGTAAAAG |
| helix64-helix72 | GATTAGTAATAACATCACTTGGTTTTTTGGGGTCGAGGTGCCGTA |
| helix65-helix67 | AGTCTGTCCATCACGCAAATTAATGCTTTCCTCGTTAGAATCAGAGC |
| helix66-helix59 | CCAGAATCCTGAGAAGTGTTTTTATAGGCCAACAGAGATAGAACCCTTCT |
| helix67-helix68 | GGGAGCTAAACAGGAGGCCGATTCCGCTACAGGGCGCGTACTATGGT |
| helix68-helix70 | TGCTTTGACGAGCACGTATAACGGAGAAAGGAAGGGAAGAAAGCGAA |
| helix69-helix49 | CCACCACACCCGCCGCGCTTAATGCGAACATTATCATTTTGCGGAACAAA |
| helix70-helix44 | AGGAGCGGGCGCTAGGGCGCTGGTACCTTTTACATCGGGAGAAACAATAA |
| helix71-helix64 | CGGGGAAAGCCGGCGAACGTGGCCCGTTGTAGCAATACTTCTTT |
| helix72-helix74 | AAGCACTAAATCGGAACCCTAAAAACAAGAGTCCACTATTAAAGAAC |
| helix73-helix87 | ACTACGTGAACCATCACCCAAATCAATTTAAATATGCAACTAAAGTACGG |
| helix74-helix76 | GTGGACTCCAACGTCAAAGGGCGTCCTGTTTGATGGTGGTTCCGAAA |
| helix75-helix43 | TTGAGTGTTGTTCCAGTTTGGCAAGTTACAAAATCGCGCAGAGGC |
| helix76-helix38 | TCGGCAAAATCCCTTATAAATCAAGACGCTGAGAAGAGTCAATA |
| helix77-helix85 | TGGTTTGCCCCAGCAGGCGAAAATAGATACATTTCGCAAATGGTCAATAA |
| helix78-helix34 | GCCCTTCACCGCCTGGCCCTGAGAATGGTTTGAAATACCGACCG |
| helix79-helix81 | GCCAGGGTGGTTTTTCTTTTCACACTCACATTAATTGCGTTGCGCTC |
| helix80-helix30 | AATGAATCGGCCAACGCGCGGGGCAGAGGCATTTTCGAGCCA |
| helix81-helix107 | ACTGCCCGCTTTCCAGTCGGGAACTGATAAATTGTGTCGAAATCCGCGAC |
| helix82-helix84 | TGGGGTGCCTAATGAGTGAGCTAGCGCGAGCTGAAAAGGTGGCATCA |
| helix83-helix91 | ACAATTCCACACAACATACGACAAAAAGATTAAGAGGAAGCCCGA |
| helix84-helix86 | ATTCTACTAATAGTAGTAGCATTAACAGTTGATTCCCAATTCTGCGA |
| helix85-helix78 | CCTGTTTAGCTATATTTTCATTTGGGCAGTGAGACGGGCAACAGCTGATT |
| helix86-helix73 | ACGAGTAGATTTAGTTTGACCATAAAAACCGTCTATCAGGGCGATGGCCC |
| helix87-helix89 | TGTCTGGAAGTTTCATTCCATATAGGATTAGAGAGTACCTTTAATTG |
| helix88-helix63 | GAATATAATGCTGTAGCTCAACATGTCCTGAGTAGAAGAACTCAAACTAT |
| helix89-helix90 | CTCCTTTTGATAAGAGGTCATTTATTCGAGCTTCAAAGCGAACCAGA |
| helix90-helix83 | CCGGAAGCAAACTCCAACAGGTCGTGTGAAATTGTTATCCGCTC |
| helix91-helix93 | AAGACTTCAAATATCGCGTTTTACCCTCAAATGCTTTAAACAGTTCA |
| helix92-helix106 | ATAGTCAGAAGCAAAGCGGATTGCATAGGCGCAGACGGTCAATCATAAGG |
| helix93-helix95 | GAAAACGAGAATGACCATAAATCAAGAAGTTTTGCCAGAGGGGGTAA |
| helix94-helix61 | TCATAAATATTCATTGAATCCACCTACATTTTGACGCTCAAT |
| helix95-helix97 | TAGTAAAATGTTTAGACTGGATATAACGCCAAAAGGAATTACGAGGC |
| helix96-helix104 | ACCAAAATAGCGAGAGGCTTTTGCAACTTCATCAAGAGTAATCTTGACAA |
| helix97-helix99 | ATAGTAAGAGCAACACTATCATATAAAACGAACTAACGGAACAACAT |
| helix98-helix60 | CATTCAACTAATGCAGATACAATTGGCAGATTCACCAGTCACACG |
| helix99-helix55 | TATTACAGGTAGAAAGATTCATCCTGCAACAGTGCCACGCTGAGAGCCAG |
| helix100-helix102 | TTGGGAAGAAAAATCTACGTTAAAGAAACACCAGAACGAGTAGTAAA |
| helix101-helix120 | ACCTTATGCGATTTTAAGAACCCTGTAGCATTCCACAGACAGCCC |
| helix102-helix116 | TTGGGCTTGAGATGGTTTAATTTCTCCAAAAAAAAGGCTCCAAA |
| helix103-helix96 | AGTGAATAAGGCTTGCCCTGACGACCCTCGTTTACCAGACGACGATAAAA |
| helix104-helix112 | GAACCGGATATTCATTACCCAAACACCCTCAGCAGCGAAAGACA |
| helix105-helix92 | ACCAGGCGCATAGGCTGGCTGACAAAAATCAGGTCTTTACCCTGACTATT |
| helix106-helix108 | GAACCGAACTGACCAACTTTGAAATTATACCAAGCGCGAAACAAAGT |
| helix107-helix82 | CTGCTCCATGTTACTTAGCCGGAACGGCCGGAAGCATAAAGTGTAAAGCC |
| helix108-helix109 | ACAACGGAGATTTGTATCATCGCAAAGAGGCAAAAGAATACACTAAA |
| helix109-helix111 | ACACTCATCTTTGACCCCCAGCGCGGCTACAGAGGCTTTGAGGACTA |
| helix110-helix29 | CACTACGAAGGCACCAACCTAAAACGCCGACAAAAGGTAAAGTAATTCTG |
| helix111-helix113 | AAGACTTTTTCATGAGGAAGTTTGGTCGCTGAGGCTTGCAGGGAGTT |
| helix112-helix105 | GCATCGGAACGAGGGTAGCAAAGAGGACAGATGAACGGTGTACAG |
| helix113-helix115 | AAAGGCCGCTTTTGCGGGATCGTTTTATCAGCTTGCTTTCGAGGTGA |
| helix114-helix27 | ATCGCCCACGCATAACCGATATATTCCCTGAACAAGAAAAATAATATCCC |
| helix115-helix117 | ATTTCTTAAACAGCTTGATACCGGGAACAACTAAAGGAATTGCGAAT |
| helix116-helix103 | AGGAGCCTTTAATTGTATCGGTCAACGTAACAAAGCTGCTCATTC |
| helix117-helix118 | AATAATTTTTTCACGTTGAAAATATTTTGCTAAACAACTTTCAACAG |
| helix118-helix22 | TTTCAGCGGAGTGAGAATAGAAAGCGAACCTCCCGACTTGCGGGAGGTTT |
| helix119-helix101 | AGTAAATGAATTTTCTGTATGGGCAACTTTAATCATTGTGAATT |
| helix120-helix122 | TCATAGTTAGCGTAACGATCTAAGCCACCCTCAGAGCCACCACCCTC |
| helix121-helix54 | TTTCGTCACCAGTACAAACTACAACGCCTTGCTGAACCTCAAATATCAAA |
| helix122-helix124 | ATTTTCAGGGATAGCAAGCCCAAGGGTTGATATAAGTATAGCCCGGA |
| helix123-helix21 | AGAACCGCCACCCTCAGAACCTATTTTGCACCCAGCTACAATTTT |
| helix124-helix17 | ATAGGTGTATCACCGTACTCAGGGGAAGCGCATTAGACGGGAGA |
| helix125-helix52 | CAGGCGGATAAGTGCCGTCGAGACTTTAGGAGCACTAACAACTAATAGAT |
| helix126-helix128 | AGACTCCTCAAGAGAAGGATTAGTAATAAGTTTTAACGGGGTCAGTG |
| helix127-helix48 | ATTTCGGAACCTATTATTCTGTATCATCATATTCCTGATTATCAG |
| helix128-helix130 | CCTTGAGTAACAGTGCCCGTATAAAATAAATCCTCATTAAAGCCAGA |
| helix129-helix16 | CTTTTGATGATACAGGAGTGTACTGGAAGTCAGAGGGTAATTGAGCGCTA |
| helix130-helix132 | ATGGAAAGCGCAGTCTCTGAATTGAGCCGCCACCAGAACCACCACCA |
| helix131-helix46 | TTGGCCTTGATATTCACAAACTACCATATCAAAATTATTTGC |
| helix132-helix40 | GAGCCGCCGCCAGCATTGACAGGTAAATCAATATATGTGAGTGAATAACC |

Figure S2: Staples sequences for synthesis of nanosphere.

| name (5'-3') | sequence |
| --- | --- |
| staple 1 | TCAGGCTGCGCAACCTAGGGCGCTGGCAATCGTCTGAAATGG |
| staple 2 | CATAACGCCAAAAGTTGCTAAACAACTTCCAATAGGAACCCA |
| staple 3 | CCGCTTCTGGTGCCCCACACCCGCCGCGACAGGAAAAACGCT |
| staple 4 | TATCGGCCTCAGGAATGGTTGCTTTGACTTGCTGGTAATATC |
| staple 5 | CATCGTAACCGTGCGAATCAGAGCGGGAATAACATCACTTGC |
| staple 6 | GATTGACCGTAATGTTAGACAGGAACGGTCACGCAAATTAAC |
| staple 7 | TCAGTTGAGATTTAAAGGAACAACTAAACCACCCTCAGAGCC |
| staple 8 | AACGAACTAACGGATGAAAATCTCCAAAGGTTTAGTACCGCC |
| staple 9 | TATACCAGTCAGGAGTATCGGTTTATCAATATAAGTATAGCC |
| staple 10 | ATCATTGTGAATTAAGCTTGATACCGATTTTTGCTCAGTACC |
| staple 11 | CGAGTAGTAAATTGGCCCACGCATAACCAGAGGCTGAGACTC |
| staple 12 | TCATTCAGTGAATAGAGTTAAAGGCCGCTGCCTATTTCGGAA |
| staple 13 | AGAACCGGATATTCAAAGACAGCATCGGGTGCCTTGAGTAAC |
| staple 14 | GGCGCATAGGCTGGTTGAGGACTAAAGAGATGATACAGGAGT |
| staple 15 | TGACCAACTTTGAAGGGTAAAATACGTATCTCTGAATTTACC |
| staple 16 | GCCGGAACGAGGCGCGAAAGAGGCAAAACAAACAAATAAATC |
| staple 17 | GATAAATTGTGTCGCCCAGCGATTATACGAAGTATGTTGAGG |
| staple 18 | TTGCGTATTGGGCGCTTTTCACCAGTGAAATAGATTAGAGCC |
| staple 19 | TATCATAACCCTCGCGTCTTTCCAGACGGTACAAACTACAAC |
| staple 20 | CAGCTGCATTAATGGCCTGGCCCTGAGATGAGGAAGGTTATC |
| staple 21 | TTGCGCTCACTGCCTGCCCCAGCAGGCGATCAATATCTGGTC |
| staple 22 | GCCTGGGGTGCCTATCGGCAAAATCCCTTCTAAAGCATCACC |
| staple 23 | ACAATTCCACACAAGGTTGAGTGTTGTTCCTGCAACAGTGCC |
| staple 24 | TCATGGTCATAGCTAGAACGTGGACTCCGCAGAAGATAAAAC |
| staple 25 | TCGACTCTAGAGGAAGGGCGATGGCCCAAGCCCTAAAACATC |
| staple 26 | GTTGTAAAACGACGTTTTGGGGTCGAGGAATATTTTTGAATG |
| staple 27 | GGCGATTAAGTTGGAAGGGAGCCCCCGAGAACCCTTCTGACC |
| staple 28 | CTTCGCTATTACGCACGTGGCGAGAAAGACACGACCAGTAAT |
| staple 29 | GAGAAGTGTTTTTAGTCGGATTCTCCGTAAATGTGAGCGAGT |
| staple 30 | ATTATTTACATTGGAATTAATTACATTTCGTTATACAAATTC |
| staple 31 | CATGGAAATACCTAATGGAAACAGTACACGGAATCATAATTA |
| staple 32 | CAGAACAATATTACTGCTTCTGTAAATCACCGACCGTGTGAT |
| staple 33 | CTGAGTAGAAGAACTCCTTGAAAACATATAGTTAATTTCATC |
| staple 34 | CGTTGTAGCAATACGAGTCAATAGTGAATCGCAAGACAAAGA |
| staple 35 | GAGGCCACCGAGTAACCTTTTTAACCTCGTTGGGTTATATAA |
| staple 36 | ACCACCCTCATTTTCAAAGACAAAAGGGAACAAAGTTACCAG |
| staple 37 | ACCCTCAGAACCGCGTAAATATTGACGGATCTTACCGAAGCC |
| staple 38 | CGGAATAGGTGTATCGTCACCGACTTGAAGCCCAATAATAAG |
| staple 39 | AGGCGGATAAGTGCCACCAGTAGCACCATGAGCGCTAATATC |
| staple 40 | CTCAAGAGAAGGATCAATGAAACCATCGGGGAGAATTAACTG |
| staple 41 | CCTATTATTCTGAAAATCAAGTTTGCCTTTTACAGAGAGAAT |
| staple 42 | AGTGCCCGTATAAACGGCATTTTCGGTCGAAACGATTTTTTG |
| staple 43 | GTACTGGTAATAAGTTTCATAATCAAAATTACAAAATAAACA |
| staple 44 | GTTCCAGTAAGCGTCGCCTCCCTCAGAGTACCAACGCTAACG |
| staple 45 | GCCTGTAGCATTCCCAACATATAAAAGAGCAGTATGTTAGCA |
| staple 46 | CTCATTAAAGCCAGAGCCACCACCCTCATAGTTGCTATTTTG |
| staple 47 | CAGGTCAGACGATTCGCCGCCAGCATTGACCTCCCGACTTGC |
| staple 48 | GTCAATAGATAATACAACTCGTATTAAAAAGGCTTATCCGGT |
| staple 49 | TAAAATATCTTTAGAAGTTTGAGTAACAAGGAATCATTACCG |
| staple 50 | AGTTGGCAAATCAACAGAAGGAGCGGAACGCACTCATCGAGA |
| staple 51 | TTGCTGAACCTCAAATGGCAATTCATCAGTCTTTCCTTATCA |
| staple 52 | ACGCTGAGAGCCAGTCTGAATAATGGAAATCCTAATTTACGA |
| staple 53 | AGAGGTGAGGCGGTTTGCACGTAAAACATATCAACAATAGAT |
| staple 54 | GCCATTAAAAATACGTTTAACGTCAGATGACAATAAACAACA |
| staple 55 | GCTATTAGTCTTTACGGGAGAAACAATAATAAAGTACCGACA |
| staple 56 | TGTACCGTAACACTTTTTGTCACAATCAGGAATACCCAAAAG |
| staple 57 | TGAAAGCGTAAGAAAAGTTACAAAATCGGTAATTTAGGCAGA |
| staple 58 | AAAAGGGACATTCTCCTGAGCAAAAGAATAGGGCTTAATTGA |
| staple 59 | TTACCAGTATAAAGCGGTAATCGTAAAAATCGGTGCGGGCCT |
| staple 60 | CTAGAAAAAGCCTGTGATAATCAGAAAAGCGCCATTCGCCAT |
| staple 61 | AAATAAGGCGTTAAAAATATTTAAATTGCCAGCTTTCCGGCA |
| staple 62 | TTCTGACCTAAATTATTAAATTTTTGTTGGGGACGACGACAG |
| staple 63 | ACGCGAGAAAACTTACGCCATCAAAAATGGTGTAGATGGGCG |
| staple 64 | AAGGAAACCGAGGACGTCATAAATATTCAACTAATGCAGATA |
| staple 65 | CTATATGTAAATGCCTTTCATCAACATTGGGAACAAACGGCG |
| staple 66 | CTTTTTAAGAAAAGGTTCAGAAAACGAGGGTAGAAAGATTCA |
| staple 67 | AGCAAGAAACAATGTACCCTGACTATTAATCTACGTTAATAA |
| staple 68 | AGAGAGATAACCCAAAAGATTAAGAGGAAAGAACTGGCTCAT |
| staple 69 | AACACCCTGAACAATAATTCGAGCTTCATAATTTCAACTTTA |
| staple 70 | AACATAAAAACAGGACAGGTCAGGATTAGAGAAACACCAGAA |
| staple 71 | TTTAACGTCAAAAAAAGAGGTCATTTTTCGTAACAAAGCTGC |
| staple 72 | GCCATATTATTTATTATAATGCTGTAGCGAGTAATCTTGACA |
| staple 73 | AGCGTCTTTCCAGATACGGTGTCTGGAACGGTGTACAGACCA |
| staple 74 | CACCCAGCTACAATAATTCTGCGAACGATAAGGGAACCGAAC |
| staple 75 | GGGAGGTTTTGAAGTTCGCAAATGGTCACTCCATGTTACTTA |
| staple 76 | ATTCTAAGAACGCGGGGCGCGAGCTGAATTGTATCATCGCCT |
| staple 77 | CGCCCAATAGCAAGTAGCATTACATCCACGGGGAGAGGCGGT |
| staple 78 | ACAAGCAAGCCGTTAGCAAAATTAAGCAGAAACCTGTCGTGC |
| staple 79 | TTCCAAGAACGGGTGGTTGTACCAAAAATCACATTAATTGCG |
| staple 80 | GCATGTAGAAACCAAGAAGCCTTTATTTGCATAAAGTGTAAA |
| staple 81 | AACTGGCATGATTATAGTAAAATGTTTAAGTAAGAGCAACAC |
| staple 82 | AAGTCCTGAACAAGCTCATATATTTTAAATTGTTATCCGCTC |
| staple 83 | TGTTCAGCTAATGCAGATTCAAAAGGGTGCTCGAATTCGTAA |
| staple 84 | AAAGGTAAAGTAATATCAATATGATATTTGCATGCCTGCAGG |
| staple 85 | GGCATTTTCGAGCCGGAGAGGGTAGCTATTCCCAGTCACGAC |
| staple 86 | GAATCGCCATATTTGTCATTGCCTGAGAGGGATGTGCTGCAA |
| staple 87 | CTGAGGCTTGCAGGAGGCTTGCCCTGACGAGAGTACCTTTAA |
| staple 88 | AAATCGGAACCCTAGTAACGCCAGGGTTTTTTTGAGAGATCT |
| staple 89 | ATATGTACCCCGGTTTTAGTATCATATGAACAATTTCATTTG |
| staple 90 | AAGATTGTATAAGCATAAGAATAAACACTAAATCAATATATG |
| staple 91 | TTGTTAAAATTCGCTAATGGTTTGAAATGTCGCTATTAATTA |
| staple 92 | TTTAACCAATAGGATTTCAAATATATTTGCGATAGCTTAGAT |
| staple 93 | CTTCCTGTAGCCAGTGATGCAAATCCAATTTATCAAAATCAT |
| staple 94 | AAATGCTTTAAACATAAGCAGATAGCCGCGACATTCAACCGA |
| staple 95 | AAAAATCAGGTCTTAAATAGCAATAGCTAAATTATTCATTAA |
| staple 96 | GCGGATTGCATCAACAAGAATTGAGTTAGCCATTTGGGAATT |
| staple 97 | GGAAGCAAACTCCAGAAGCGCATTAGACATAGCAGCACCGTA |
| staple 98 | TTGCTCCTTTTGATTGAAAATAGCAGCCTTAGCGTCAGACTG |
| staple 99 | GCTTAATTGCTGAACCCAATCCAAATAAATAGCCCCCTTATT |
| staple 100 | ATATGCAACTAAAGGCCTAATTTGCCAGTCACCGGAACCAGA |
| staple 101 | AACAGTTGATTCCCTTTATCCTGAATCTCCGCCACCCTCAGA |
| staple 102 | GCCAGAGGGGGTAAAGACTCCTTATTACAACGCAAAGACACC |
| staple 103 | TATATTTTCATTTGAGGCGTTTTAGCGAACAGGAGTAGACTT |
| staple 104 | TCTACTAATAGTAGCAAATCAGATATAGTCCTTTGCCCGAAC |
| staple 105 | CAAGGCAAAGAATTTTTATTTTCATCGTTTATCATTTTGCGG |
| staple 106 | CATAAAGCTAAATCATTAAACCAAGTACTTATCATCATATTC |
| staple 107 | AATACTTTTGCGGGATCAATAATCGGCTATATAATCCTGATT |
| staple 108 | TAATGTGTAGGTAAAGAACGCGCCTGTTGAAATAAAGAAATT |
| staple 109 | ACAGTCAAATCACCTCTGTCCAGACGACGAATATACAGTAAC |
| staple 110 | GATAAATTAATGCCAGTAATAAGAGAATACGGATTCGCCTGA |
| staple 111 | CAATACTGCGGAATAACGCAATAATAACATAGAAAATTCATA |
| staple 112 | ACAAAGGCTATCAGAACAACGCCAACATCGCAGAGGCGAATT |
| staple 113 | AGAGAATCGATGAACCAACGCTCAACAGGATGATGAAACAAA |
| staple 114 | GAAAGGAGCGGGCGTGTTGGGAAGGGCGCTAGCATGTCAATC |
| staple 115 | ACAGGGCGCGTACTAGATCGCACTCCAGTAAACGTTAATATT |
| staple 116 | CGATTAAAGGGATTGGATAGGTCACGTTAATTCGCGTCTGGC |
| staple 117 | TAATTTTTTCACGTACAACATTATTACAAATGACCATAAATC |
| staple 118 | TGAATTTCTTAAACCCTTATGCGATTTTAGCCCGAAAGACTT |
| staple 119 | CGGCTACAGAGGCTCTGACCTTCATCAATCAACATGTTTTAA |
| staple 120 | GCACCAACCTAAAACAGACGGTCAATCAGTAGATTTAGTTTG |
| staple 121 | AGTACAAGGTTTTTCCAGGGTCGGAGATAAGGTGGCATCAAT |
| staple 122 | GGTCCACGCTGGTTCGCTTTCCAGTCGGATAAAGCCTCAGAG |
| staple 123 | ATAGCCCGAGATAGCATACGAGCCGGAACAACGCAAGGATAA |
| staple 124 | TTCTGTATGGGATTGAATTACGAGGCATGACTGGATAGCGTC |
| staple 125 | AAAAACCGTCTATCTCCCCGGGTACCGAGAGAAAGGCCGGAG |
| staple 126 | AGCGAGAGGCTTTTATAAAAACCAAAAT |
| staple 127 | TAGTTAGCGTAACGACAGACAGCCCTCA |
| staple 128 | ATACATAAAGGTGGAACGTAGAAAATAC |
| staple 129 | CAAATATCGCGTTTAGTCAGAGGGTAATTTACCATTAGCAAG |
| staple 130 | ACCATTAGATACATCCTTAAATCAAGATGAGCCGCCACCAGA |
| staple 131 | AAATTTTTAGAACCAAAAATAATATCCCGGGTTAGAACCTAC |
| staple 132 | GCTGCGCGTAACCAGGAAACCAGGCAAAGCCCCAAAAACAGG |
| staple 133 | TGCTTTCCTCGTTAATCTGCCAGTTTGAAAATCAGCTCATTT |
| staple 134 | GGAGTGAGAATAGAGGAATACCACATTCATTGAATCCCCCTC |
| staple 135 | AGGAGCCTTTAATTCGTTGGGAAGAAAATAGTCAGAAGCAAA |
| staple 136 | TGACAACAACCATCGGCTTGAGATGGTTAAGCGAACCAGACC |
| staple 137 | ATCTAAAGTTTTGTTTTACCAGACGACGGCAAAAGAAGTTTT |
| staple 138 | CACCCTCAGCAGCGATTACCCAAATCAAGCGGATGGCTTAGA |
| staple 139 | AGTTTCCATTAAACAGAGGACAGATGAAGTTTCATTCCATAT |
| staple 140 | ACTCATCTTTGACCAAATCCGCGACCTGATAACCTGTTTAGC |
| staple 141 | GATTGCCCTTCACCAATCGGCCAACGCGATAAATCATACAGG |
| staple 142 | TGGTGGTTCCGAAAATGAGTGAGCTAACCATTATGACCCTGT |
| staple 143 | GAGTCCACTATTAAGTTTCCTGTGTGAAATGCAATGCCTGAG |
| staple 144 | ACCCAAATCAAGTTGCCAGTGCCAAGCTCAACCGTTCTAGCT |
| staple 145 | GGGAAAGCCGGCGACAGCTGGCGAAAGGGTCTGGAGCAAACA |
| staple 146 | ACGGAATAAGTTTAGAGTTTCGTCACCATTAGTAAATGAATT |
| staple 147 | TGGTTTACCAGCGCCAGGGATAGCAAGCTCAACAGTTTCAGC |
| staple 148 | TTGAGGGAGGGAAGCACCCTCAGAACCGGGAATTGCGAATAA |
| staple 149 | AGGTGAATTATCACCACCGTACTCAGGAAAAAAGGCTCCAAA |
| staple 150 | AGAGCCAGCAAAATCGTCGAGAGGGTTGGCTTGCTTTCGAGG |
| staple 151 | GCCGGAAACGTCACTAGGATTAGCGGGGAGTTGCGCCGACAA |
| staple 152 | ATCAGTAGCGACAGACATGAAAGTATTAGATATATTCGGTCG |
| staple 153 | TAGCGCGTTTTCATCAGTTAATGCCCCCTTTTGCGGGATCGT |
| staple 154 | AGCGTTTGCCATCTTTTTAACGGGGTCAAACGAGGGTAGCAA |
| staple 155 | GCCACCACCGGAACCATACATGGCTTTTCTTTTTCATGAGGA |
| staple 156 | ACCGCCACCCTCAGAATGGAAAGCGCAGATGCCACTACGAAG |
| staple 157 | ACCACCACCAGAGCGGCCTTGATATTCAGAATACACTAAAAC |
| staple 158 | TACAAACAATTCGACATTTGAGGATTTACAAGCGCGAAACAA |
| staple 159 | GTTATTAATTTTAAGAGCACTAACAACTGACGGGCAACAGCT |
| staple 160 | AACAAAGAAACCACCAGTTGAAAGGAATGAGTTGCAGCAAGC |
| staple 161 | CTGATTATCAGATGATATCAAACCCTCAAAAATCCTGTTTGA |
| staple 162 | GTTTGGATTATACTCAGCAAATGAAAAATATAAATCAAAAGA |
| staple 163 | CATATCAAAATTATCAGTATTAACACCGCCAGTTTGGAACAA |
| staple 164 | GCGTAGATTTTCAGCGAACGAACCACCAAACGTCAAAGGGCG |
| staple 165 | AGTACCTTTTACATATGCGCGAACTGATCTACGTGAACCATC |
| staple 166 | TTGCTTTGAATACCTACGTGGCACAGACTGCCGTAAAGCACT |
| staple 167 | ATTCATTTCAATTAGGCCAACAGAGATATTTAGAGCTTGACG |
| staple 168 | CATCAAGAAAACAACAGATTCACCAGTCGAAGGGAAGAAAGC |
| staple 169 | AATTACCTTTTTTACATTTTGACGCTCAAGTGTAGCGGTCAC |
| staple 170 | TGAGTGAATAACCTCGCCAGCCATTGCACTTAATGCGCCGCT |
| staple 171 | ATTTTCCCTTAGAATCAAACTATCGGCCGAGCACGTATAACG |
| staple 172 | TAAGACGCTGAGAATTCTTTGATTAGTAGCTAAACAGGAGGC |
| staple 173 | AGGTCTGAGAGACTAAAGAGTCTGTCCATACGCCAGAATCCT |

Figure S3: Staples sequences for synthesis of nanorod.

#### Relative capacitance

The oscillator frequency can be assumed linear with respect to the loading capacitance $C_{IN}$ when the change in the capacitance recorded by the sensor is sufficiently small as shown in Eq. 1.

$$f(C_{IN}) = -\alpha.(C_{IN} + C_0) + f_0 \tag{1}$$

Here, $\alpha$ is the pixel sensitivity, $C_0$ is the parasitic capacitance and $f_0$ is the baseline frequency as discussed in.[3]

To eliminate the need for calibration and find out $C_0$ and $f_0$ for each pixel, we take the difference of the responses with respect to *no deposition* state of the sensor (i.e., *idle* as the sample)

$$f(C_{idle}) = -\alpha.(C_{idle} + C_0) + f_0 \tag{2}$$

Hence, for any *sample* deposited on the sensor (i.e., buffer, nanostructure, etc.) we get a similar equation as

$$f(C_{sample}) = -\alpha.(C_{sample} + C_0) + f_0 \tag{3}$$

Subtraction of Eq. 2 from Eq. 3 and rearranging,

$$f(C_{sample}) = -\alpha.(C_{sample} - C_{idle}) + f(C_{idle}) \tag{4}$$

Defining $\Delta C_{sample} = C_{sample} - C_{idle}$, Eq. 4 can be written as

$$\Delta C_{sample} = \frac{-(f(C_{sample}) - f(C_{idle}))}{\alpha} \tag{5}$$

Now, the effect of the nanostructure on the capacitance is the difference between the capacitive change due to the deposited sample and the effect of the buffer on the capacitance.

$$\Delta C_{nanostructure} = C_{sample} - C_{buffer}$$

While $f(C_{sample})$, $f(C_{idle})$ and $f(C_{buffer})$ are available from sensor readout, $\alpha$ needs to be predetermined with the calibration process.[3] However, since the sensitivity coefficient is temperature dependent, it varies with the ambient conditions of the day. Therefore, we seek to obtain a normalized metric that is independent of $\alpha$. Hence, we introduce a relative capacitance metric, (*relative* $C$ or $r\Delta C$) as defined below. We calculate $r\Delta C$ as percentage of change with respect to the reference buffer.

$$r\Delta C = 100 \times \left| \frac{\Delta C_{nanostructure}}{\Delta C_{buffer}} \right|$$

Substituting from Eq-1,

$$r\Delta C = 100 \times \left| \frac{-(f(C_{sample}) - f(C_{buffer}))}{\alpha} \times \frac{-\alpha}{f(C_{buffer}) - f(C_{idle})} \right|$$

$$r\Delta C = 100 \times \left| \frac{(f(C_{sample}) - f(C_{buffer}))}{f(C_{buffer}) - f(C_{idle})} \right|$$

Rearranging the terms, we get,

$$\boxed{r\Delta C = 100 \times \left| 1 - \frac{f(C_{sample}) - f(C_{idle})}{f(C_{buffer}) - f(C_{idle})} \right|} \tag{6}$$

Note that, the normalization of the $\Delta C_{sample}$ can be performed with any choice of reference sample. Using the buffer as a reference sample incorporates the negative control group for the nanostructure study in the relative capacitance metric as explained in the Results section. For our $Mg^{2+}$ studies in Figure 4, we calculate the $r\Delta C$ of 1xTAE with varying concentrations of $MgCl_2$ by referencing it against 1xTAE with no $Mg^{2+}$.

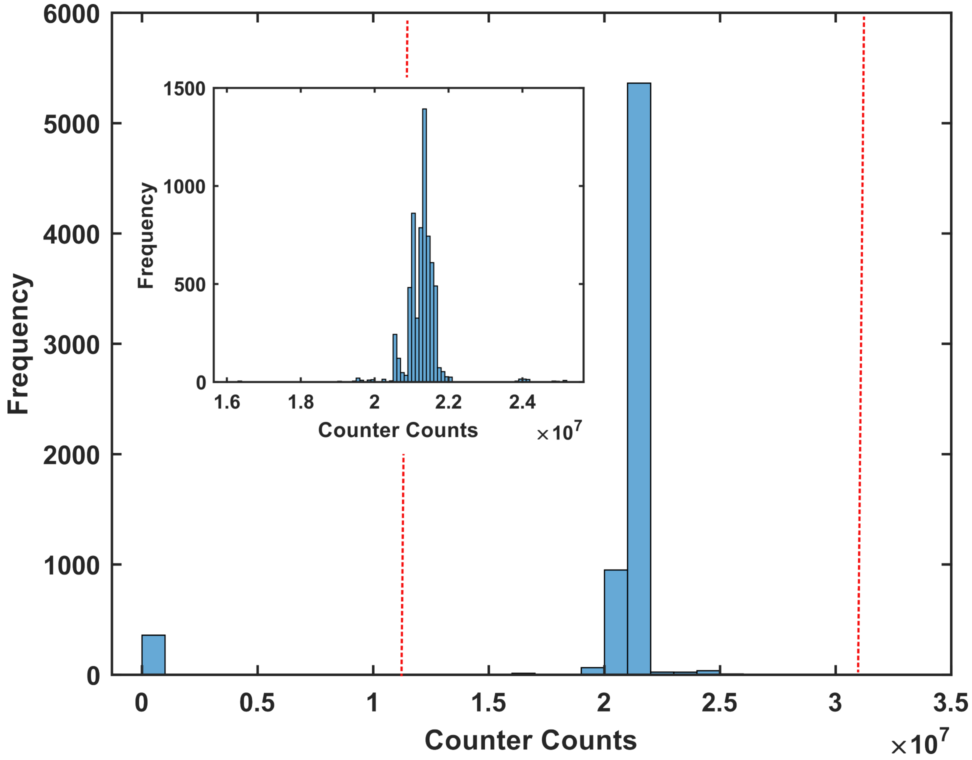


Figure S4: Data histogram before and after (inset) glitch correction: Histogram is plotted to visually evaluate the raw counts and set thresholds (here we used 50%) values above and below the median count to eliminate random glitches (zeros or very high values) resulting from non-ideal readout circuit.

*Temporal noise filtering:* Once the erroneous data points are removed (histogram in the inset of Fig. S4), we fit the remaining data with Savitzky-Golay filter to smooth out the temporal noise and discover the data trend as shown in Fig. S5.

After that we take the median of each trend signal from each pixel and average them to get the sample point for a particular trial (the dashed line in Fig. S6).

#### Data post-processing

**Underlying assumptions.** The frequency of the oscillator, $f(C_{sample})$ for a particular sample can be calculated as

$$f(C_{sample}) = \frac{N_{T_s}}{T_s} \tag{7}$$

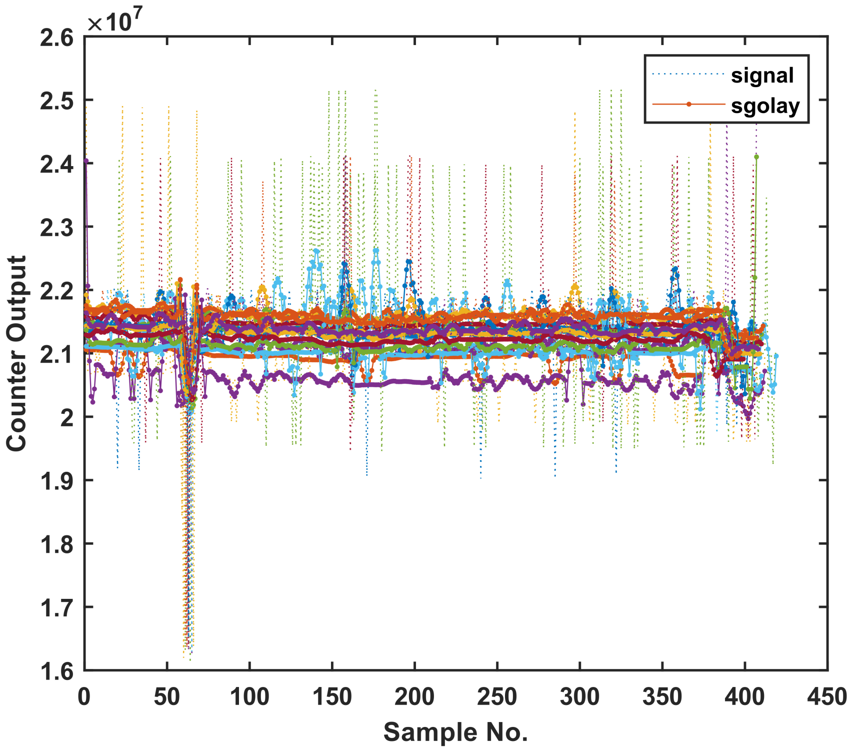


Figure S5: Application of Savitsky-Golay smoothing filter: To extract the data trend from each pixel we applied a third order Savitsky-Golay filter with a frame length of 11 on the glitch corrected data. The dotted lines shows the raw counts and the solid lines show the smoothed data trend. The filter minimizes the effect of the temporal noise from data point extraction.

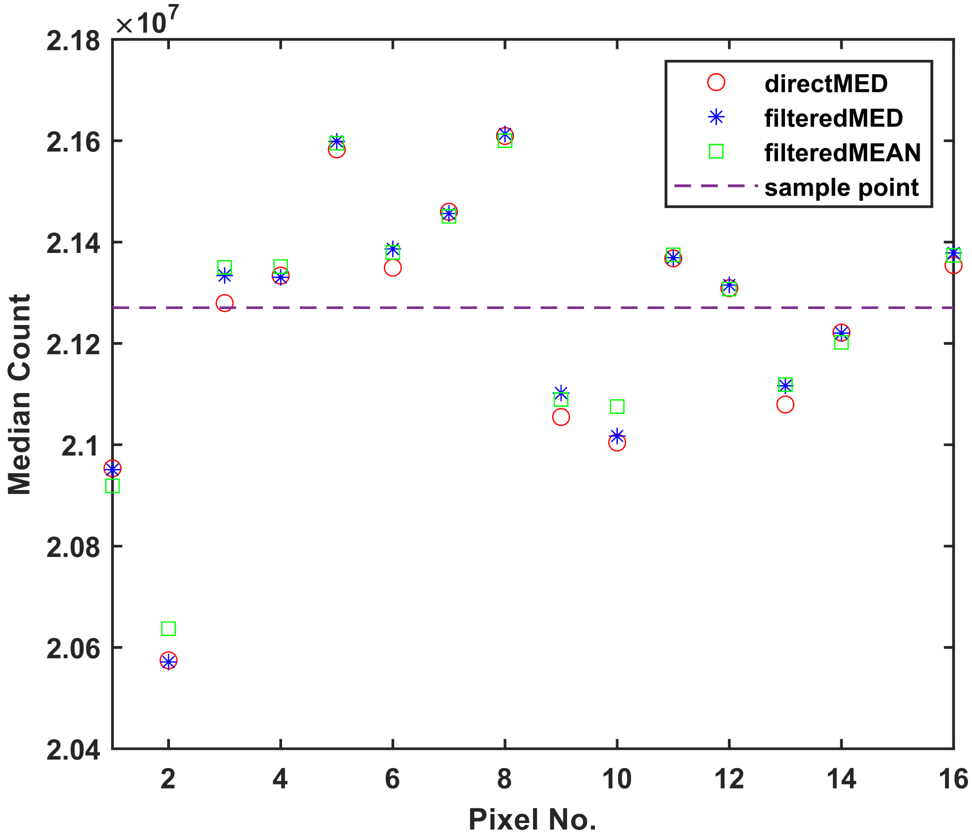


Figure S6: Data point extraction: The final data point for a particular sample is extracted by taking the average of the median counts from each pixel after applying the smoothing filter (dashed line). The direct median counts from the raw data are comparable to the filtered median (and filtered mean) counts, implying that for a system with low computational power might skip the temporal filtering and yet get a reasonably accurate measure.

Here, $N_{T_s}$ is the number of oscillations registered with in the sampling time $T_s$. If the sampling time is kept constant for all the terms involved in Eq. 6, we can use the oscillator counts directly as the raw data. In this work, the sampling time was kept constant (30 s) for all data collection. An alternative method is to calculate the frequencies explicitly before processing to noise cancellation.

**Random glitch correction.** We assume that capacitive response of our sensor to deposition, such as a solution containing DNA origami nanostructures, is not instantaneous. Therefore, we consider sporadic large values or zeros to be associated with non-ideal sensor readout noise. To eliminate the noise outliers, we use thresholding to discard the data points outside the 50% from the median in the data histogram as illustrated in Fig. S4.

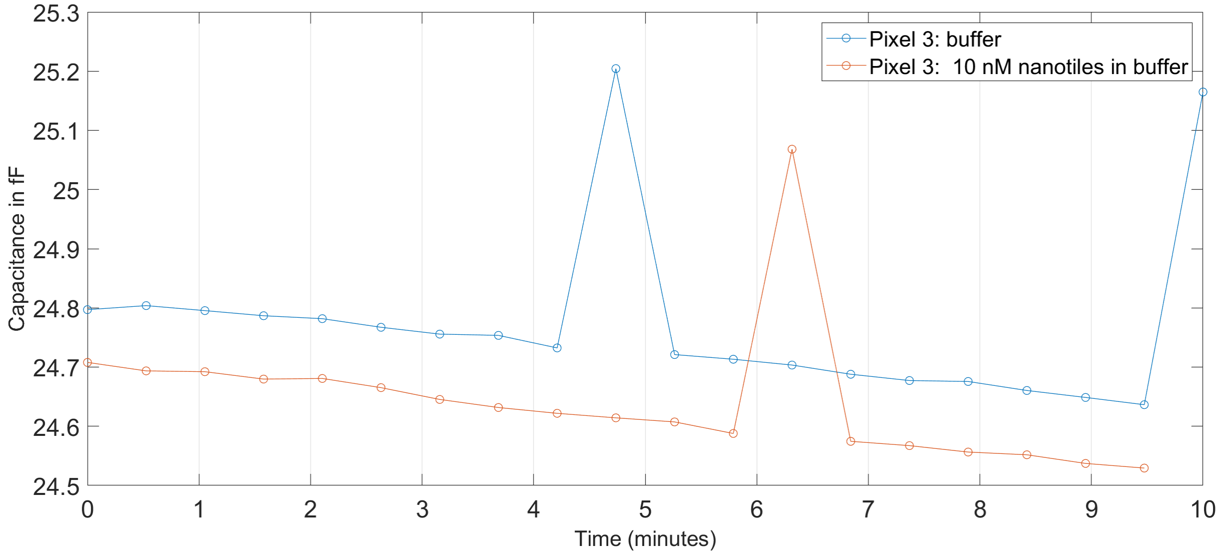


Figure S7: For a randomly selected pixel, Pixel-3, the estimated capacitance for buffer, 1xTAE with 12.5 mM $Mg^{2+}$, and 10 nM nanotiles in 1xTAE with 12.5 $Mg^{2+}$ are shown as a time series of 10 minutes measured on the same day. For this plot, we assume $|\alpha|$=590 kHertz/fF, clock frequency = 4e6 kHertz to compute estimated capacitance from the ring oscillator counts.[3]

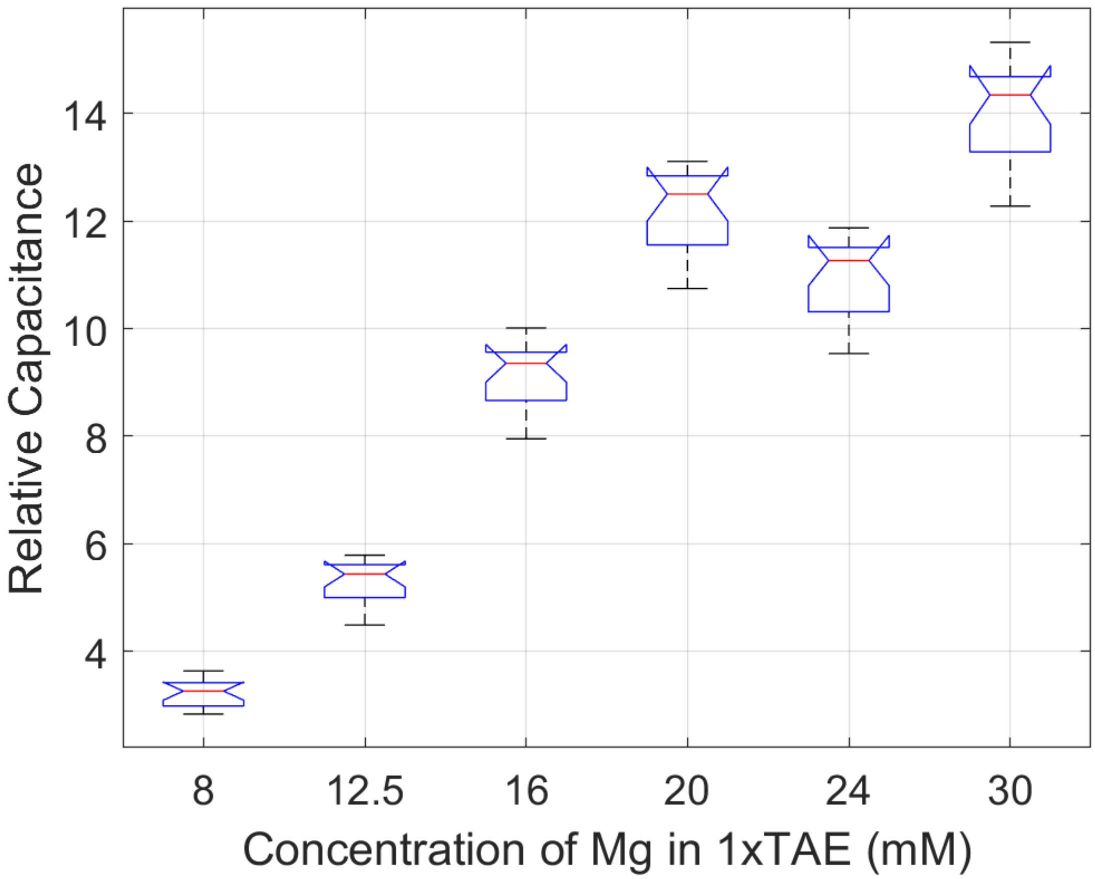


Figure S8: The box plots of relative capacitance as $Mg^{2+}$ is varied in 1xTAE. The notch (red bar) on the box plot represents the median over all pixels and corresponds to the scatter plot in Figure-4a.

.

Table S1: Tukey-Kramer pair-wise comparison for 1xTAE $Mg^{2+}$ sweep (N=3-4, $p < 0.01$).

| Buffer A $Mg^{2+}$ (mM) in 1xTAE | Buffer B $Mg^{2+}$ (mM) in 1xTAE | $\Delta = \mu_A - \mu_B$ | Confidence Interval for $\Delta$ |
|---|---|---|---|
| 8 | 12.5 | -2.057 | (-2.727, -1.388) |
| 12.5 | 16 | -3.867 | (-4.536, -3.197 ) |
| 16 | 20 | -3.0377 | (-3.708, -2.368 ) |
| 16 | 24 | -1.806 | (-2.476, -1.137 ) |
| 20 | 24 | 1.231 | (0.561, 1.901 ) |
| 20 | 30 | -1.857 | (-2.526, -1.187 ) |
| 24 | 30 | -3.088 | (-3.757, -2.418 ) |

Table S2: Tukey-Kramer pair-wise comparison for 10 nM pre-annealed nanotiles with excess staples ratio (K) (N=3, $p < 0.01$).

| Excess ratio A | Excess ratio B | $\Delta = \mu_{K,A} - \mu_{K,B}$ | Confidence Interval for $\Delta$ |
|---|---|---|---|
| 1 | 5 | -0.545 | (-0.731,-0.358) |
| 1 | 10 | -1.322 | (-1.509,-1.135) |
| 5 | 10 | -0.778 | (-0.964,-0.591) |

Table S3: Tukey-Kramer pair-wise comparison for 10 nM DNA origami nanotile formation (N=3, $p < 0.01$).

| Nanostructure Concentration (nM) | $\Delta = \mu_{preannealed} - \mu_{nanotiles}$ | Confidence Interval for $\Delta$ |
|---|---|---|
| 5 | -0.614 | (-0.764, -0.463) |
| 10 | -0.412 | (-0.561, -0.262) |
| 15 | -0.512 | (-0.662, -0.362) |
| 20 | 0.077 | (-0.073, 0.227) |

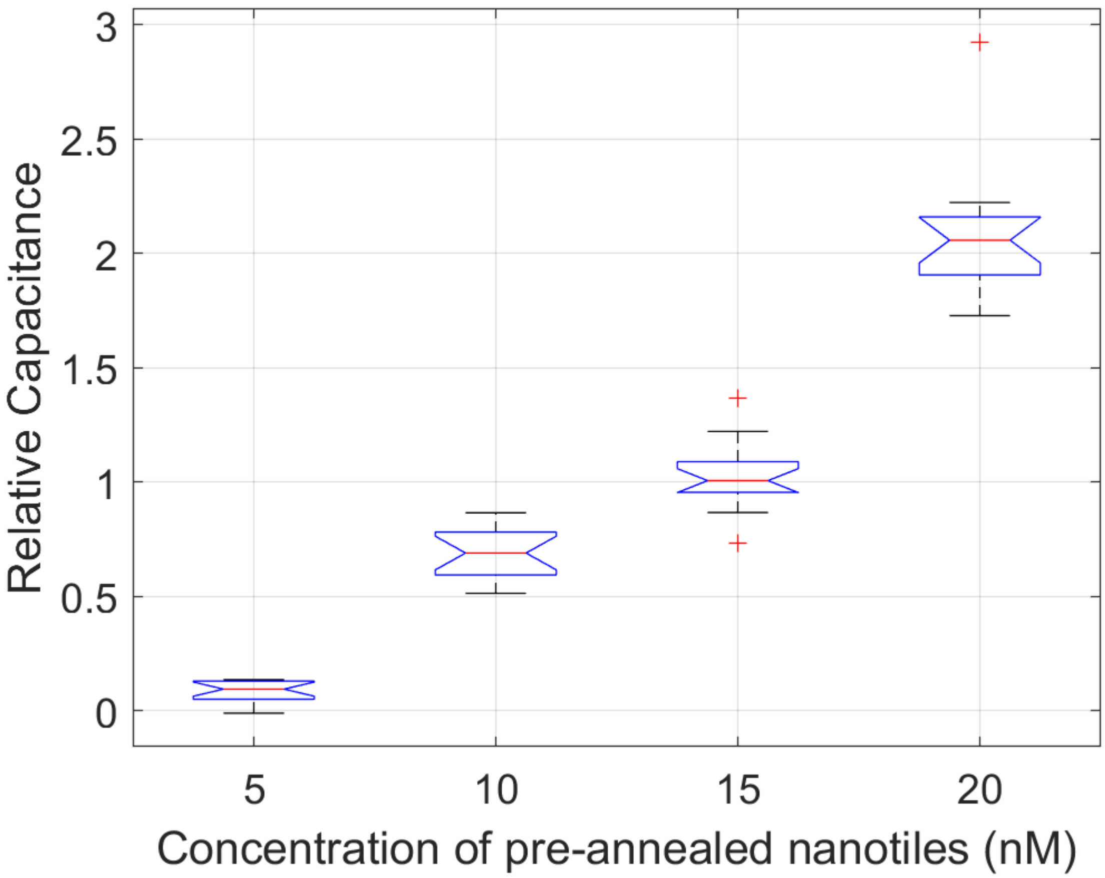


Figure S9: The box plots of relative capacitance as concentration of pre-annealed nanotiles is varied in from 5 nM to 20 nM in 1xTAE with 12.5 mM $Mg^{2+}$. The notch (red bar) on the box plot represents the median over all pixels and corresponds to the scatter plot for pre-annealed nanotiles in Figure-5(b).

.

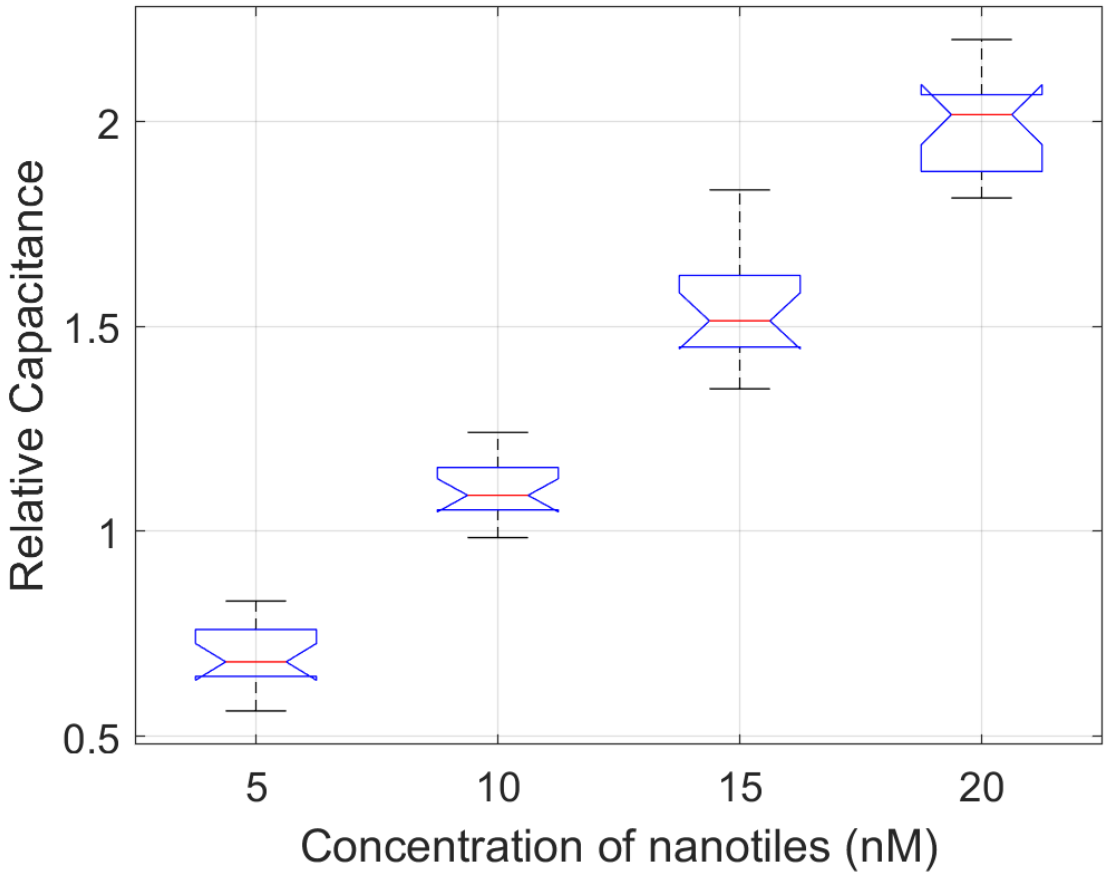


Figure S10: The box plots of relative capacitance as concentration of nanotiles is varied in from 5 nM to 20 nM in 1xTAE with 12.5 mM $Mg^{2+}$. The notch (red bar) on the box plot represents the median over all pixels and corresponds to the scatter plot for nanotiles in Figure-5(b) and Figure-6(a).

.

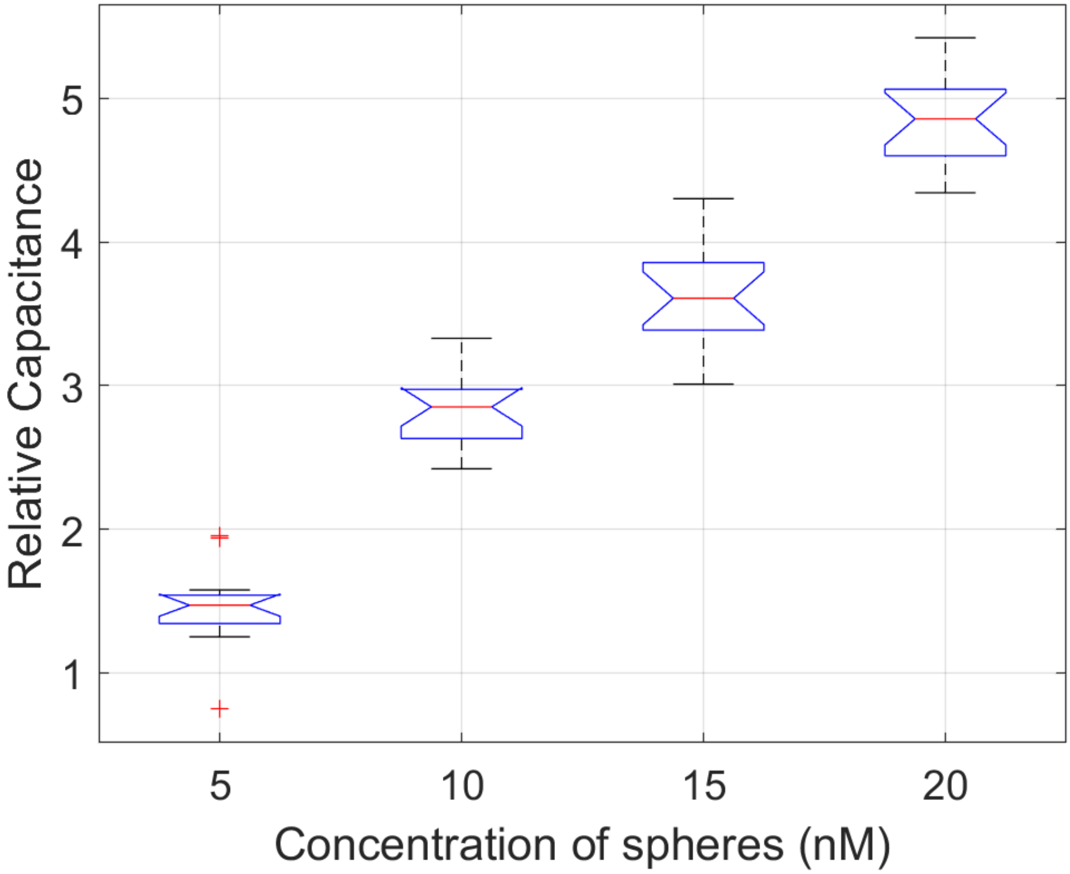


Figure S11: The box plots of relative capacitance as concentration of nanospheres is varied in from 5 nM to 20 nM in 1xTAE with 12.5 mM $Mg^{2+}$. The notch (red bar) on the box plot represents the median over all pixels and corresponds to the scatter plot for nanospheres in Figure-6(a).

.

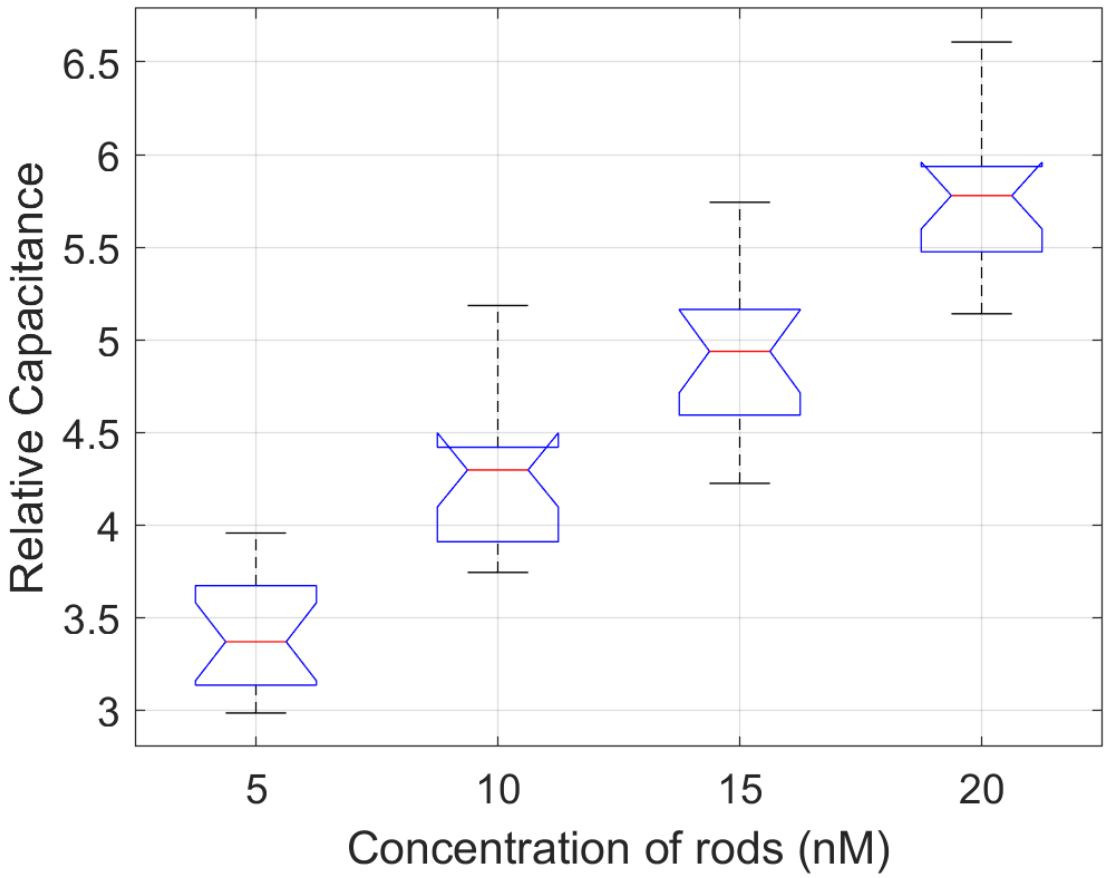


Figure S12: The box plots of relative capacitance as concentration of nanorods is varied in from 5 nM to 20 nM in 1xTAE with 12.5 mM $Mg^{2+}$. The notch (red bar) on the box plot represents the median over all pixels and corresponds to the scatter plot for nanorods in Figure-6(a).

.

Table S4: Tukey-Kramer pair-wise comparison for nanotiles and nanospheres (N=3, $p < 0.01$).

| Nanostructure Concentration (nM) | $\Delta = \mu_{nanotiles}$ - $\mu_{nanospheres}$ | Confidence Interval for $\Delta$ |
|---|---|---|
| 5 | -0.755 | (-1.085, -0.425) |
| 10 | -1.729 | (-2.059, -1.4) |
| 15 | -2.096 | (-2.426, -1.767) |
| 20 | -2.834 | (-3.164, -2.504) |

Table S5: Tukey-Kramer pair-wise comparison for nanotiles and nanorods (N=3, $p < 0.01$).

| Nanostructure Concentration (nM) | $\Delta = \mu_{nanotiles}$ - $\mu_{nanorods}$ | Confidence Interval for $\Delta$ |
|---|---|---|
| 5 | -2.706 | (-3.036,-2.376) |
| 10 | -3.174 | (-3.504,-2.844) |
| 15 | -3.394 | (-3.724,-3.065) |
| 20 | -3.745 | (-4.075,-3.415) |

Table S6: Tukey-Kramer pair-wise comparison for nanospheres and nanorods (N=3, $p < 0.01$).

| Nanostructure Concentration (nM) | $\Delta = \mu_{nanospheres}$ - $\mu_{nanorods}$ | Confidence Interval for $\Delta$ |
|---|---|---|
| 5 | -1.951 | (-2.281,-1.626) |
| 10 | -1.445 | (-1.774,-1.115) |
| 15 | -1.298 | (-1.628,-0.968) |
| 20 | -0.911 | (-1.241,-0.581) |

### References

(1) Liu, Y.; Wijesekara, P.; Kumar, S.; Wang, W.; Ren, X.; Taylor, R. E. The effects of overhang placement and multivalency on cell labeling by DNA origami. *13*, 6819–6828.

(2) Wang, W.; Hayes, P. R.; Ren, X.; Taylor, R. E. Synthetic cell armor made of DNA origami. *(Submitted)*.

(3) Senevirathna, B. P.; Lu, S.; Dandin, M. P.; Basile, J.; Smela, E.; Abshire, P. A. Real-Time Measurements of Cell Proliferation Using a Lab-on-CMOS Capacitance Sensor Array. *12*, 510–520.
